# Plasma proteome analyses in individuals of European and African ancestry identify *cis*-pQTLs and models for proteome-wide association studies

**DOI:** 10.1101/2021.03.15.435533

**Authors:** Jingning Zhang, Diptavo Dutta, Anna Köttgen, Adrienne Tin, Pascal Schlosser, Morgan E. Grams, Benjamin Harvey, CKDGen Consortium, Bing Yu, Eric Boerwinkle, Josef Coresh, Nilanjan Chatterjee

## Abstract

Improved understanding of genetic regulation of proteome can facilitate the identification of causal mechanisms for complex traits. We analyzed data on 4,657 plasma proteins from 7,213 European American (EA) and 1,871 African American (AA) individuals from the ARIC study, and further replicated findings on 467 AA individuals from the AASK study. Here we identified 2,004 proteins in EA and 1,618 in AA, with majority overlapping, which showed associations with common variants in *cis*-regions. Availability of AA samples led to smaller credible sets and significant number of population-specific *cis*-pQTLs. Elastic-net produced powerful models for protein prediction in both populations. An application of proteome-wide association studies (PWAS) to serum urate and gout, implicated several proteins, including *IL1RN,* revealing the promise of the drug anakinra to treat acute gout flares. Our study demonstrates the value of large and diverse ancestry study for genetic mechanisms of molecular phenotypes and their relationship with complex traits.

## Introduction

Genome-wide association studies (GWAS) to date have cumulatively mapped tens of thousands of loci containing common genetic variants associated with complex traits ^1, 2^. As the majority of the variants are in non-coding regions ^3, 4^, researchers have focused on understanding the role of gene-expression regulation as a mechanism for complex trait genetic association ^5–9^ . In the future, comprehensive understanding of causal mechanisms for complex traits will require the integration of data from various types of genomic and molecular traits ^10^. Proteins, the ultimate product of the transcripts, are subject to post-translational modifications and processing, and contain additional information that cannot be detected at the level of the transcriptome.

Recently, major opportunities have arisen to substantially increase our understanding of the causal role of proteins in complex traits due to availability of an accurate high throughput technology for measuring proteins in different types of samples ^11, 12^. The plasma proteome has received particular attention as it can capture a wide variety of proteins that are active in different biological processes ^13^. The proteome is often dysregulated by diseases, and it is highly amenable for drug targeting ^14, 15^. A number of genetic studies have identified protein quantitative trait loci (pQTL), for plasma ^14–19^ as well as some other tissues ^20–22^, and noted that pQTLs are enriched for GWAS associations across an array of complex traits ^14–22^. Studies have used pQTLs as instruments in conducting Mendelian randomization (MR) analysis to identify causative proteins, and hence potential therapeutic targets, across diverse phenotypes ^23–25^.

In spite of substantial progress, understanding of the genetic architecture of the proteome and its overlap with those of gene expressions and complex traits remains limited. While the sample size for some studies of the plasma proteome has involved thousands of individuals, it is likely that identification of pQTLs remains incomplete, both due to inadequate sample size or/and lack of comprehensive protein measurements. Further, existing proteomic studies have been mostly restricted to samples of European ancestry, and thus cannot inform potential heterogeneity by ancestry. Additionally, advanced tools for incorporating pQTL information for exploring causal effects of proteins, such as those available for analysis of gene-expression ^26, 27^, are lacking.

In this article, we report results from a comprehensive set of analyses of *cis*-genetic regulation of the plasma proteome in the large European and African American cohorts of the Atherosclerosis Risk in Communities (ARIC) study ^28^. We focus on the identification of *cis*-associations, which compared to *trans*-, have been shown to more replicable across different proteomic platforms ^29^ and are less likely to be affected by horizontal pleiotropy that could pose additional challenge for downstream Mendelian-randomization analyses ^30^. We carry out a set of association and fine-mapping analyses to identify common (minor allele frequency (MAF) > 1%) *cis*-pQTLs and compare results across ancestries to explore shared and unique genetic architecture. For each population, we characterize *cis*-heritability of the proteome due to common variants and build models for genetically predicting levels of plasma proteins. Using these models, we then conduct proteome-wide association studies (PWAS) of serum urate ^31^, an important biomarker of purine metabolism with high heritability and large available large GWAS summary statistics, and the complex disease gout, which can result from high urate levels ^31^. We create several data resources for using our results to inform future studies (http://nilanjanchatterjeelab.org/pwas).

## Results

### Identification of cis-pQTLs Across Two Populations

We performed separate *cis*-pQTL analyses for the African American (AA) and European American (EA) populations in the ARIC study, with total sample sizes of n = 1,871 and 7,213, respectively. We performed analyses based on plasma samples collected during the third visit of the cohort ^28^ (see Supplementary Table 1 for sample characteristics). Relative concentrations of plasma proteins or protein complexes were measured by modified aptamers (‘SOMAmer reagents’, hereafter referred to as SOMAmers) ^11, 12^.

After quality control (see Methods), we analyzed 4,657 SOMAmers, which tagged proteins or protein complexes encoded by 4,435 genes, and 204 of them were tagged by more than one SOMAmer. We defined *cis*-regions to be +/- 500Kb of the transcription start site (TSS) in the *cis*-pQTL analysis. In the *cis*-regions, we analyzed 10,961,088 common (MAF>1%) single-nucleotide polymorphisms (SNPs) for AA and 6,181,856 for EA with imputed or genotyped data after quality filtering (see Methods). For identification of *cis*-pQTLs, we performed regression analyses of protein levels after residualizing by sex, age, 10 genetic principal components (PCs) and the study sites. In addition, similar to eQTL analyses ^8^, we adjusted for Probabilistic Estimation of Expression Residuals (PEER) factors ^32, 33^ to account for hidden confounders that may influence clusters of proteins. We observed that the inclusion of PEER factors substantially improved power for *cis*-pQTL studies due to reduced residual variance (Fig. 1a, Supplementary Table 2). In all subsequent analyses, protein levels measured by SOMAmers were residualized with respect to these sets of PEER factors and then normalized by quantile-quantile transformation.

**Fig. 1:**
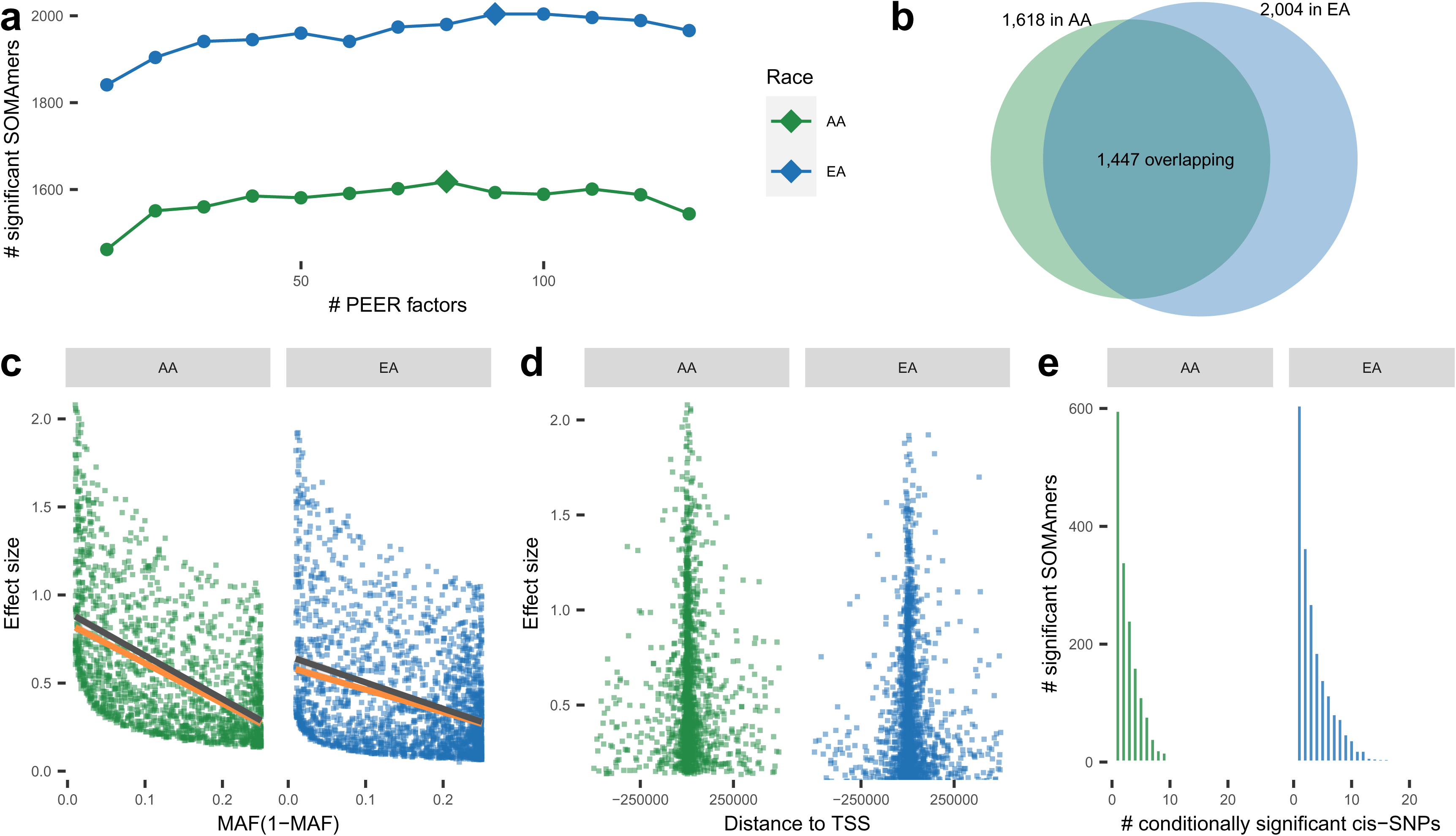
*Cis*-pQTL analysis. *Cis*-pQTL analysis overview (n = 7,213 and 1,871 for EA and AA, respectively, in ARIC). (**a**) Number of SOMAmers detected to have significant *cis*-pQTLs versus number of PEER factors used in models. Diamonds mark the numbers of PEER factors used in the following analysis which identify maximal number of significant SOMAmers. (**b**) Venn diagram of significant SOMAmers in EA and AA populations. (**c**) Effect sizes of sentinel *cis*-SNPs of pQTLs v.s. minor allele frequencies (MAF(1-MAF)). Lines are fitted with (orange) and without inverse-power weighting (dark grey). (**d**) Effect sizes of sentinel *cis*-SNPs of pQTLs v.s. distance to TSS. (**e**) Number of conditional independent *cis*-pQTLs per significant SOMAmer.

In the ARIC study, we identified a total of 2,004 and 1,618 significant SOMAmers, i.e. SOMAmer with at least one significant (at false discovery rate (FDR)<5%) *cis*-pQTL near the putative protein’s gene, in the EA and AA populations, respectively, with 1,447 of these overlapping across the populations (Fig. 1b, Supplementary Tables 3.1 and 3.2). Compared to plasma pQTL studies conducted in the past in European ancestry sample^15, 16^, we almost tripled the number of significant SOMAmers with known *cis*-pQTLs ^17, 18^ (1,465 v.s. 508 using the same Bonferroni corrected genome-wide threshold for significance) (Supplementary Table 3.1) and we successfully replicated 99% (504/508) of previously identified *cis*-pQLTs (Supplementary Table 4).

We found 10% of the sentinel *cis*-pQTLs identified in EA were non-existent or rare, defined as two or less individuals carrying the variant, in the Phase-3 1000 Genome Project (1000Genome) ^34^ African population. In contrast, nearly one third of the variants identified in the AA population were non-existent or rare in the 1000Genome European population, signifying the value of diverse ancestry data to identify ancestry-specific *cis*-pQTLs (Supplementary Tables 3.1 and 3.2). For *cis*-pQTLs which were identified through either of the two populations, but were common in (MAF>1%) in both, the effect-sizes showed high degree of concordance across the populations (Extended Data Figure 1). We further carried out a replication study using data available on additional 467 individuals from the African American Study of Kidney Disease and Hypertension (AASK) ^35^ , which also ascertained proteins using the SOMAScan platform. Among 1,398 sentinel *cis*-SNPs which were identified through the ARIC AA sample and which were genotyped or imputed in AASK, we found 93% showed effects in the same direction and 69% showed statistical significance at FDR<5% in the replication analysis (Supplementary Tables 5.1 and 5.2).

Genotypic effect sizes for *cis*-pQTLs were inversely associated with minor allele frequencies even after accounting for bias due to power for detection ^36^(Fig. 1c), and decreased with distance from the TSS (Fig. 1d). Using stepwise regression ^37, 38^, we identified multiple conditional independent *cis*-SNPs for 1,398 (70%) and 1,021 (63%) of the significant SOMAmers in EA and AA populations, respectively (Fig. 1e, Supplementary Tables 6.1 and 6.2).

Protein altering variants (PAVs) may result in apparent *cis*-pQTLs owing to altered epitope binding effects ^15^. Following a procedure recommend earlier ^15^, we found that while in the EA population, up to 65% (1,299 out of 2,004) of the sentinel pQTLs could be affected by LD with known PAVs, the corresponding proportion drops to 47% (765 out of 1,618) in the AA population (see Supplementary Tables 3.1, 3.2 and 7). However, large overlap observed between *cis*-eQTL and *cis*-pQTLs in colocalization analysis (see below) indicates they are driven by underlying causal variants and reduces concerns for any large-scale effect of epitope artifacts in the detection of *cis*-pQTLs.

### Cis-eQTL Overlap and Functional Enrichment

To evaluate the extent to which the *cis*-pQTL variants were also involved in modulating transcriptional levels, we cross referenced the *cis*-pQTLs with significant *cis*-eQTLs (at FDR<5%) from the Genotype-Tissue Expression project (GTEx) V8 ^9^ across 49 different tissues. Since the GTEx cohort is primarily of European ancestry (85.3% EA in V8), we restricted the analysis to the EA cohort only throughout the paper. We found that, approximately 73.9% of the sentinel *cis*-pQTLs, or variants in high LD (*r^2^* > 0.8) with them, were also significant *cis*-eQTLs for the same gene in at least one tissue (Extended Data Figure 2a). Further, pairwise colocalization indicated that for 49.4% of the significant SOMAmers, *cis*-pQTLs colocalize with *cis*-eQTLs in at least one of the GTEx tissues with high posterior probability (PP.H4≥80%) (Extended Data Figure 2b, Supplementary Tables 8.1 and 8.2). Further, *cis*-pQTLs tended to be significant *cis*-eQTLs across multiple tissues possibly because plasma protein level contain signatures from multiple tissues (Extended Data Figure 3).

Integrating pQTLs with the functional and regulatory annotations of the genome, curated from existing database (see Methods), offers a powerful way to understand the molecular mechanisms and consequences of genetic regulatory effects. We found that *cis*-pQTLs were enriched for several protein altering functions which may be caused by epitope binding effects noted earlier (Extended Data Figure 4a-b). After adjusting for PAVs, independent sentinel *cis*-pQTLs were enriched in a large spectrum of functional annotations including untranslated regions (5’ and 3’), promoters and transcription factor binding sites, with a pattern that was consistent across the two populations (Extended Data Figure 4c-d and Supplementary Table 9).

### Fine Mapping

To identify the causal variants underlying the significant *cis*-pQTLs for plasma proteins, we first conducted population-specific fine-mapping for the 1,447 overlapping significant SOMAmers across two populations using SuSiE ^39^ (Supplementary Tables 10.1 and 10.2). We found that the average number of variants in the 95% credible sets were significantly smaller in AA compared to that in EA (21.29 in EA v.s. 12.11 in AA; p-value = 8.43×10^-27^; Fig. 2 a-b). This is possibly driven in part by the lower average LD in AA, but also could be due to the smaller sample sizes in AA, resulting in lower statistical power. To demonstrate the added value of including two populations in identifying possibly shared causal variants, we further conducted a cross-ancestry meta-analysis using MANTRA ^40^.

**Fig. 2:**
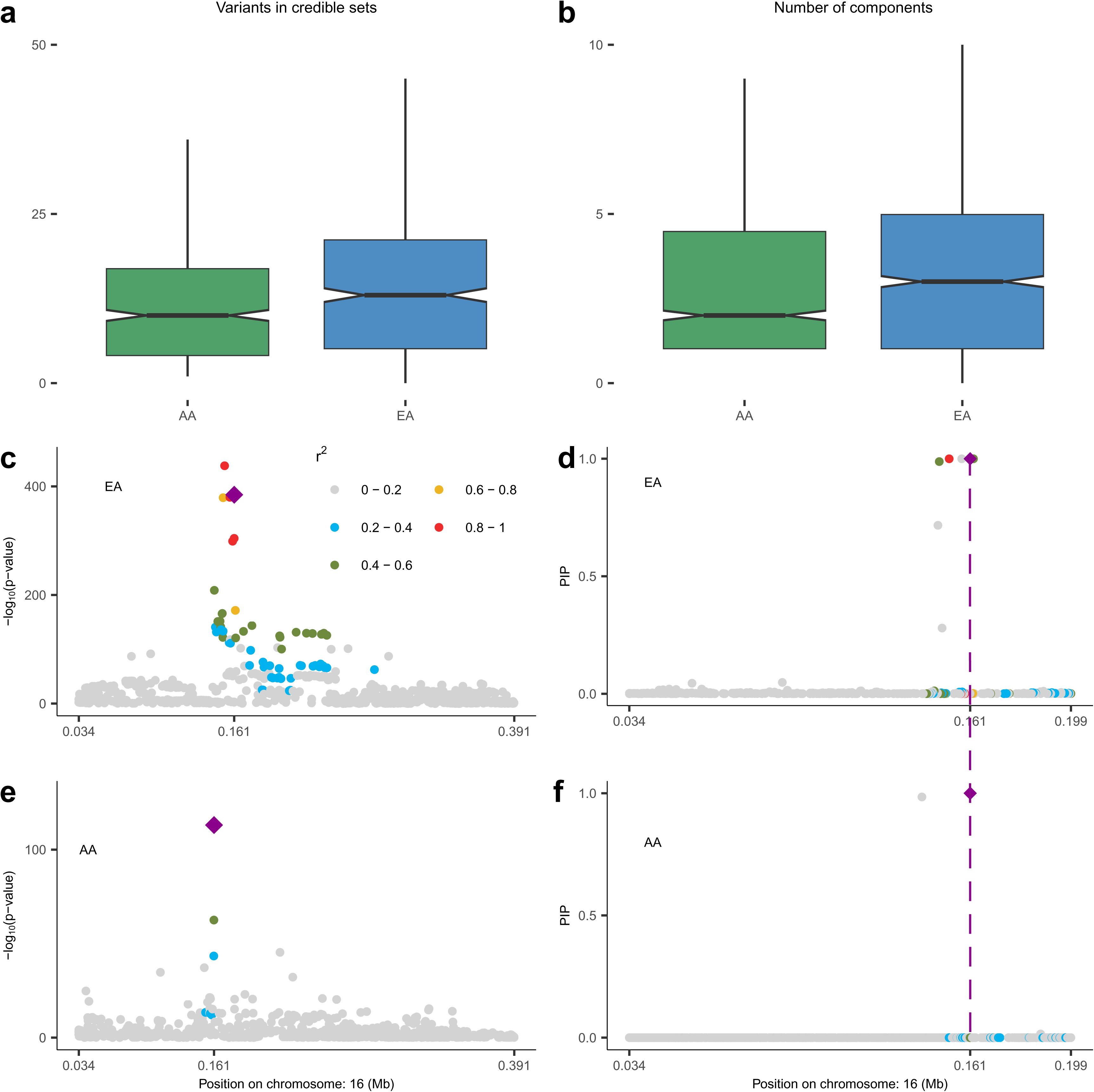
Fine-mapping analysis. (**a**) Distribution of size of credible sets and (**b**) that of number of independent SuSIE clusters across 1,447 SOMAmers that have at least one significant *cis*-pQTL in both EA and AA populations. The boxes in (**a-b**) are drawn from first and third quartiles, with the median at the center, and the whiskers extending to 1.5 times the interquartile range from the box boundaries. The power of fine-mapping using data from two populations is further illustrated using the example of *HBZ*. Regional Manhattan plots are shown based on single SNP p-value, obtained from two-sided z-test of association, and SuSIE posterior probabilities for EA (Panel **c** and **d**) and AA (Panel **e** and **f**) populations. The SNP rs2541645 (chr16: 161106; marked in diamond shape throughout) is detected as the shared causal *cis*-pQTL across the two ancestries using posterior probabilities computed by MANTRA (See Methods for more details) . The legend for the range of r^2^ between other SNPs and rs2541645 is shown at the upper right corner in (c). Sample sizes for EA and AA populations are n = 7,213 and 1,871, respectively.

As an example, we illustrate the fine-mapped *cis*-region (+/-500Kb) for *HBZ* on chromosome 16p13.3 corresponding to the Hemoglobin subunit zeta protein (HBAZ; Uniprot ID: P02008), which is involved in oxygen transport and metal-binding mechanisms ^41, 42^ and has been associated with thalassemia ^43^. After performing *cis* association analyses (Fig. 2c and 2e), fine-mapping within the EA individuals identifies a 95% credible set of seven variants (Fig. 2d) while that within the AA individuals identifies a smaller credible set of two variants only (Fig. 2f). Further, cross-ancestry meta-analysis further points to a single variant rs2541645 (16:161106 G>T) as the possible shared causal variant between the two populations. This variant was in fact the most significantly associated *cis*-pQTL for *HBZ* in AA but not in EA, and had some evidence of differences in MAF across the populations (MAF = 0.32 in EA v.s. 0.18 in AA). This SNP is a strong eQTL for *HBZ* expression in GTEx V8 whole blood (p-value = 6.7x10^-80^), and associated with several erythrocyte related outcomes in the UK Biobank including mean corpuscular hemoglobin (p-value=1.1x10^-14^) and reticulocyte fraction of red cells (p-value=3.2x10^-9^) ^44, 45^. Together, these findings suggest that rs2541645 might be a regulatory variant for *HBZ* protein levels and possibly warrant further study on downstream phenotypic consequences especially in the context of blood related mechanisms and thalassemia.

### Cis-Heritability of Proteins and Protein Imputation Models

We estimated *cis*-heritability (*cis*-h^2^) of plasma proteins, i.e. the proportion of variance of protein levels that could be explained by all *cis*-SNPs, using GCTA ^46^. We found 1,350 and 1,394 SOMAmers were *cis*-heritable, i.e., have significant non-zero *cis*-h^2^ (p-value < 0.01) (see Methods), for the EA and AA populations, respectively, and 1,109 of them overlapped (Supplementary Table 11). The majority of those significant *cis*-heritable SOMAmers also had *cis*-pQTLs identified in our study (96% for AA and 99% for EA, Supplementary Table 12). The *cis*-h^2^ for significant SOMAmers (median *cis*-h^2^ = 0.10 for AA, and 0.09 for EA) tended to be substantially smaller than those reported for gene-expression ^47^ in two related tissues ^13^, liver and whole blood, in GTEx V7 (Fig. 3a) and in GTEx V8 (Extended Data Figure 5). The pattern is expected given the closer relationship of genetic variation to transcripts than to the encoded proteins, which are subject to additional processing including post-translational modifications.

**Fig. 3:**
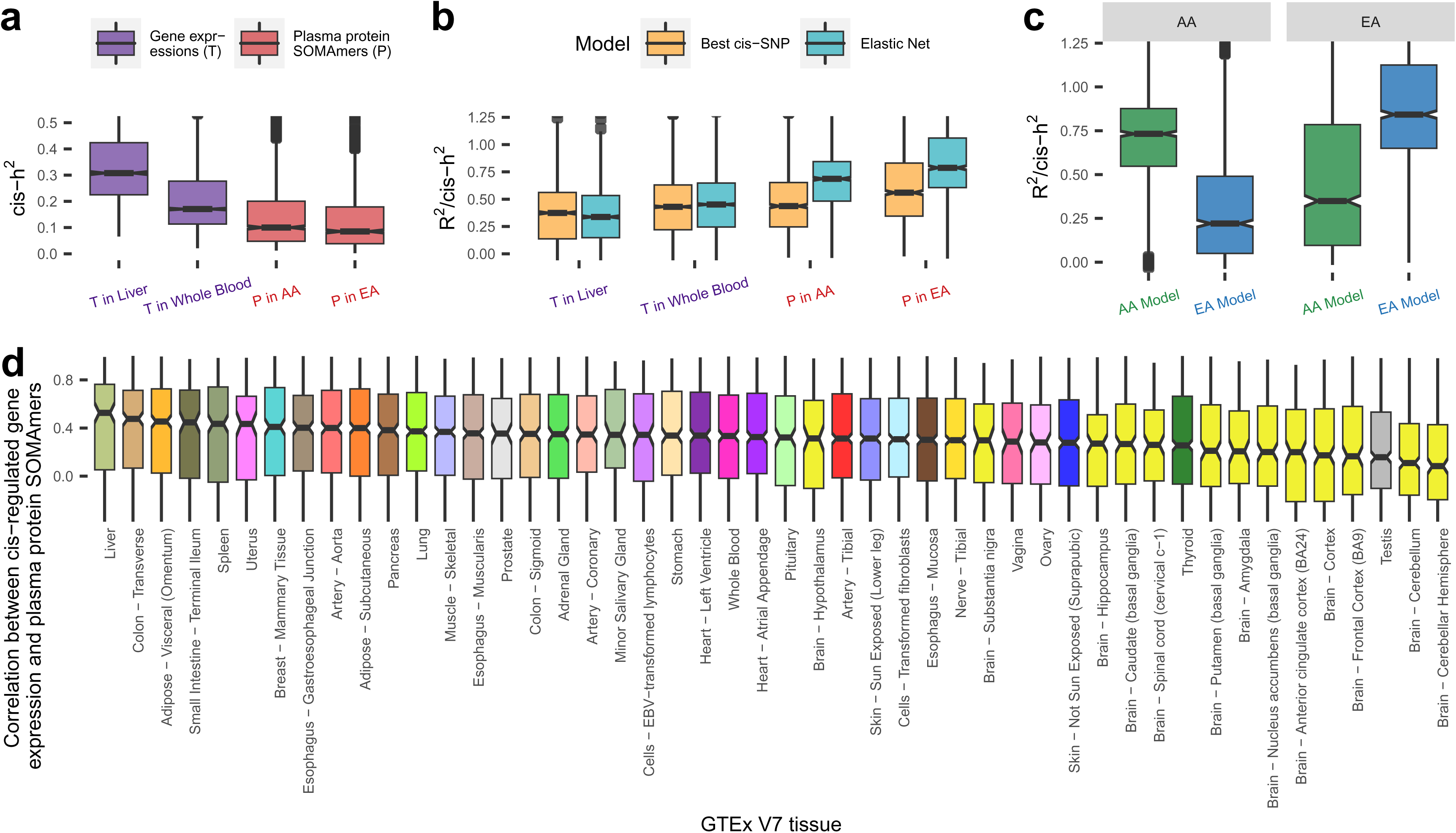
*Cis*-heritability and evaluation of models for genetic prediction of proteins. *Cis*-heritability (*cis*-h^2^) estimates and genetic imputation models are obtained using GTEx V7 data for gene expression levels, and ARIC data for plasma protein levels. Sample sizes for gene expression levels across GTEx V7 tissues are provided in Supplementary Table 13, and those for plasma protein levels in EA and AA in ARIC are n = 7,213 and 1,871, respectively. (**a**) Estimated *cis*-h^2^ for gene expression levels and plasma protein levels. (**b**) Prediction R^2^, standardized by estimated *cis*-h^2^ (R^2^/*cis*-h^2^), using imputation models trained by: the most significant *cis*-SNP; and Elastic Net using all *cis*-SNPs. (**c**) Cross-ancestry prediction accuracy by applying imputation models built from one population to the other population. (**d**) *Cis*-regulated genetic correlation between plasma proteins and expression levels for underlying genes across all GTEx (V7) tissues estimated based on 1000Genomes reference European samples (n = 498). Additional results using preliminary models available from GTEx V8 can be found in Supplementary Table 16. In boxplots, the boxes are drawn from first and third quartiles, with the median at the center, and the whiskers extending to 1.5 times the interquartile range from the box boundaries. Figures are truncated in the y-axis at *cis*-h^2^=0 and 0.5 in (a), R^2^/*cis*-h^2^=0 and 1.25 in (b-c), correlation = -0.25 and 1 in (d) for better display. *Cis*-h^2^ (a) and imputation model performances (b-d) are shown only for those gene expressions or plasma proteins which show significance *cis*-h^2^ (p-value < 0.01 in likelihood ratio test examining the significance of the random effect component in GCTA model). Exact *cis*-h^2^ estimates and p-values of their significance are provided in Supplementary Table 11 for plasma protein levels, and those for gene expression levels can be obtained from FUSION/TWAS imputation models available from http://gusevlab.org/projects/fusion/#reference-functional-data.

Next, we built protein imputation models for *cis*-heritable SOMAmers using an elastic net machine learning method as has been used for modeling gene-expression ^26^. The median accuracy for the elastic-net models for protein predictions, evaluated as the prediction R^2^ standardized by *cis-*h^2^, was 0.79 and 0.69 for the EA and AA populations, respectively. Compared with imputation models built only with the sentinel *cis*-pQTL, the elastic net models gained 36% and 40% of accuracy for the EA and AA populations, respectively (Fig. 3b, Supplementary Table 13). In cross-ancestry analysis, we found that models trained in the EA population performed worse in the AA population than the converse, in spite of a much smaller sample size in AA, again indicating the advantage of the latter population to identify causal pQTLs which are more likely to have robust effects across ancestries (Fig. 3c).

### Cis-Correlation between Plasma Proteome and Transcriptome

We then explored *cis*-regulated genetic correlation between plasma proteins and expression levels for the underlying genes across a variety of tissues. We used genotype data for Europeans from 1000Genome to evaluate Pearson’s correlation coefficients between genotypically-imputed protein levels and genotypically-imputed expression levels, with the latter being computed based on models that have been previously built and published by Gusev *et al.* ^27^ based on data from the GTEx V7 (Supplementary Tables 13 and 14). We also used models based on GTEx V8 developed by the same group (available through personal communication), but because of their preliminarily nature, we perform all main analyses using the V7 models and present preliminary results from the V8 models in supplementary data. Overall, genetically imputed plasma proteins are only moderately correlated with those for gene expression levels (Fig. 3d). Consistent with previous study ^48^, we find that plasma proteins show strongest genetic correlations with genes expression levels in the liver, the organ responsible for the synthesis of many highly abundant plasma proteins. The lowest genetic correlations were seen for brain-related tissues, which may be due to the blood-brain barrier. In GTEx V8, we observed a similar pattern for high-/low-rank tissues (Supplementary Table 15.1). The correlations between direct plasma protein measurements and imputed gene expression levels in ARIC showed similar trend but have generally lower values as they account for additional variability of protein measurements due to non-genetic factors (Extended Data Figure 6).

### Proteome-wide Association Study (PWAS) of Complex Traits

We illustrate an application of the protein imputation model by conducting proteome-wide association studies for two related complex traits: (1) serum urate, a highly heritable biomarker of health representing the end product of purine metabolism in humans, and (2) gout, a complex disease caused by urate crystal deposition in the setting of elevated urate levels and the resulting inflammatory response. We obtained GWAS summary-statistics data for these traits generated by the CKDGen Consortium ^31^ involving a total sample size of n = 288,649 and 754,056, respectively. As this GWAS was conducted primarily in EA population, we carried out the PWAS analysis using the models generated for the EA population.

We used a computational pipeline previously developed for conducting Transcriptome-wide Association Studies (TWAS) based on GWAS summary-statistics ^27, 49^ to carry out an analogous PWAS analysis. Simulation studies showed that type 1 error of PWAS analysis based on our protein imputation weights are well controlled (Extended Data Figure 7). Among all *cis*-heritable SOMEmers with imputation models, we identified 10 and 3 distinct loci containing genes for which the encoded proteins were found to be significantly (p-value < 3.7x10^-5^) associated with serum urate and gout, respectively. We further examined whether the PWAS signals could be explained by *cis*-genetic regulation of the expression of nearby (1Mb region around) genes and *vice* versa by performing bivariate analysis conditioning on imputed expression values for nearby genes that are found to be significantly associated based on the TWAS analysis. Main results were based on GTEx V7 models (Fig. 4, Table 1, Table 2, Extended Data Figure 8), and further validated using GTEx V8 preliminary models (Supplementary Table 16). For the TWAS analysis, we considered significance of genes based on two trait-relevant tissues available in GTEx V7, namely whole blood and liver, but also explored other tissues more broadly (see Methods).

**Fig. 4:**
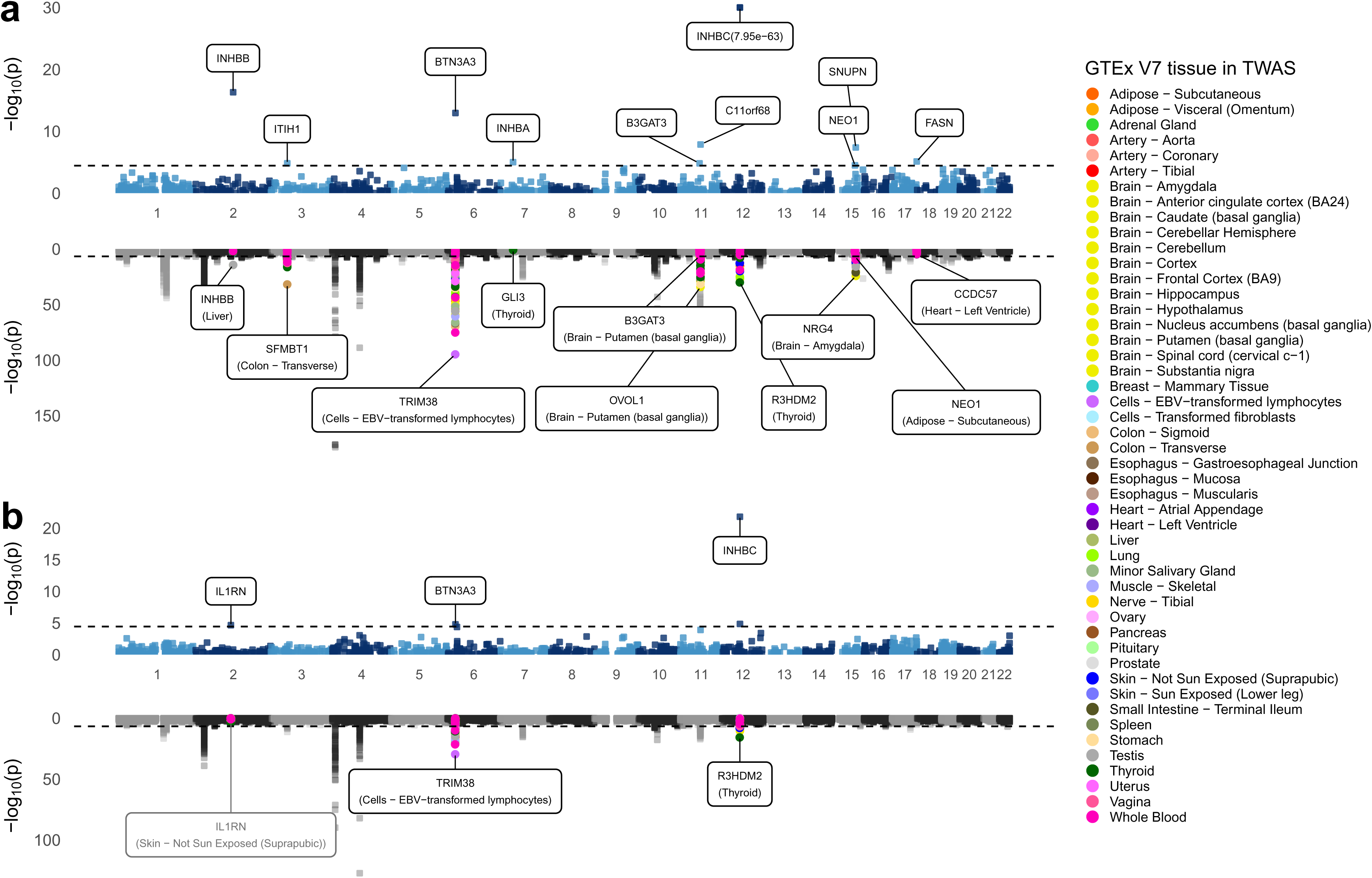
Miami plots for PWAS and TWAS analyses for serum urate level and gout. Miami plot for PWAS (upper) and TWAS (lower) of (**a**) urate and (**b**) gout. Each point represents a p-value for a two-sided z-test of association between the phenotypes and the *cis*-genetic regulated plasma protein or expression level of a gene, ordered by genomic position on the x axis and the -log10(p-value) for the association strength on the y axis. The black horizontal dash lines are the significance threshold after Bonferroni correction for the total number of imputation models (p-value = 3.7x10^-5^ for PWAS and 1.3x10^-6^ for TWAS). Urate PWAS and TWAS in (a) are truncated in the y-axis at -log_10_(p-value) = 30 and -log_10_(p-value) = 150 for better display. Nearby TWAS genes (+/- 500Kb) for significant PWAS genes are colored by GTEx tissues. The most significant nearby-TWAS gene is labelled with its gene name and corresponding tissue. The TWAS of *IL1RN* does not reach TWAS significance threshold and thereby was labeled with grey. All primary TWAS analyses are conducted based on established models developed using data from GTEx V7, and results for the identified top genes/tissue combinations are further validated using preliminary models available from GTEx V8 (Supplementary Table 16).

**Table 1.**
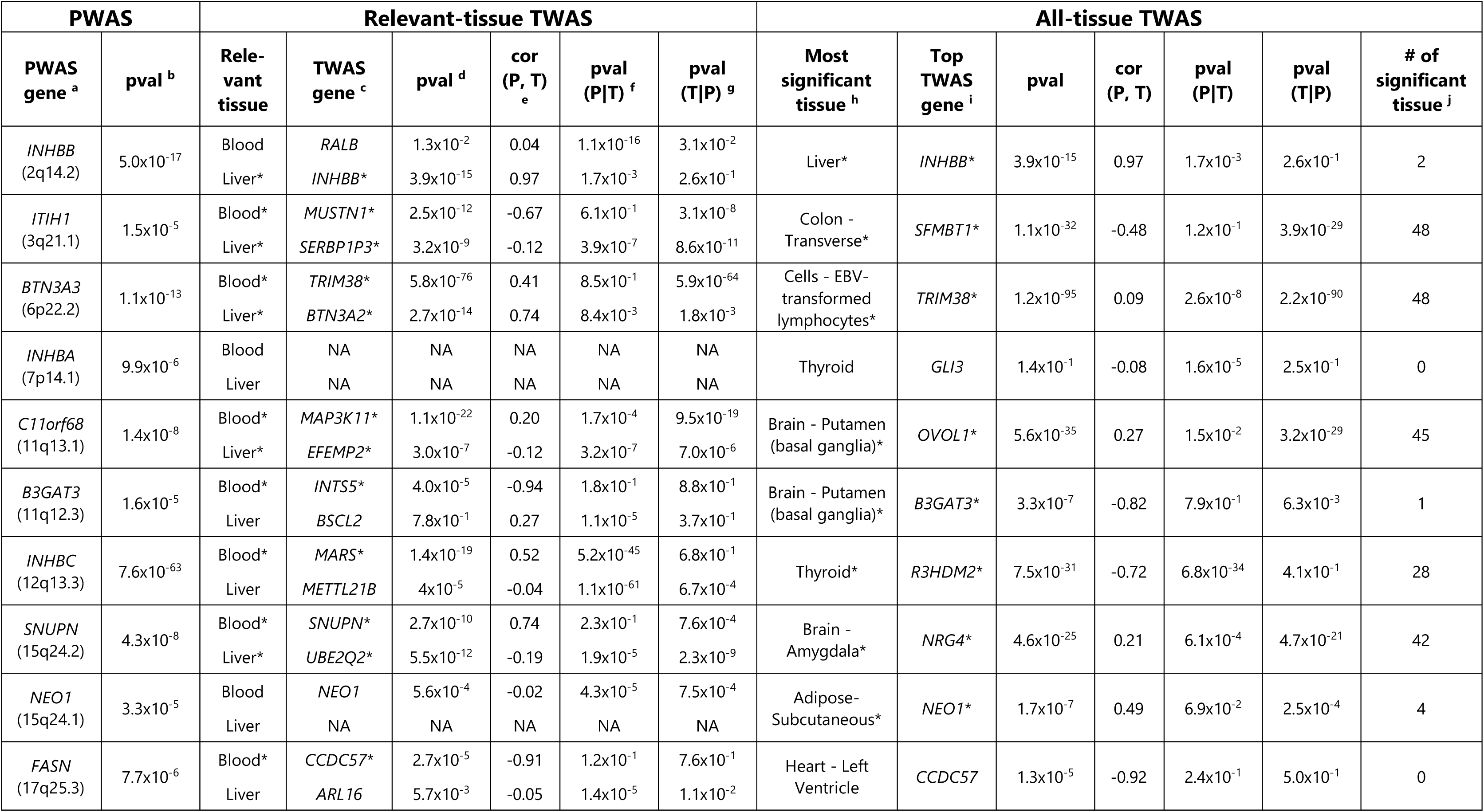
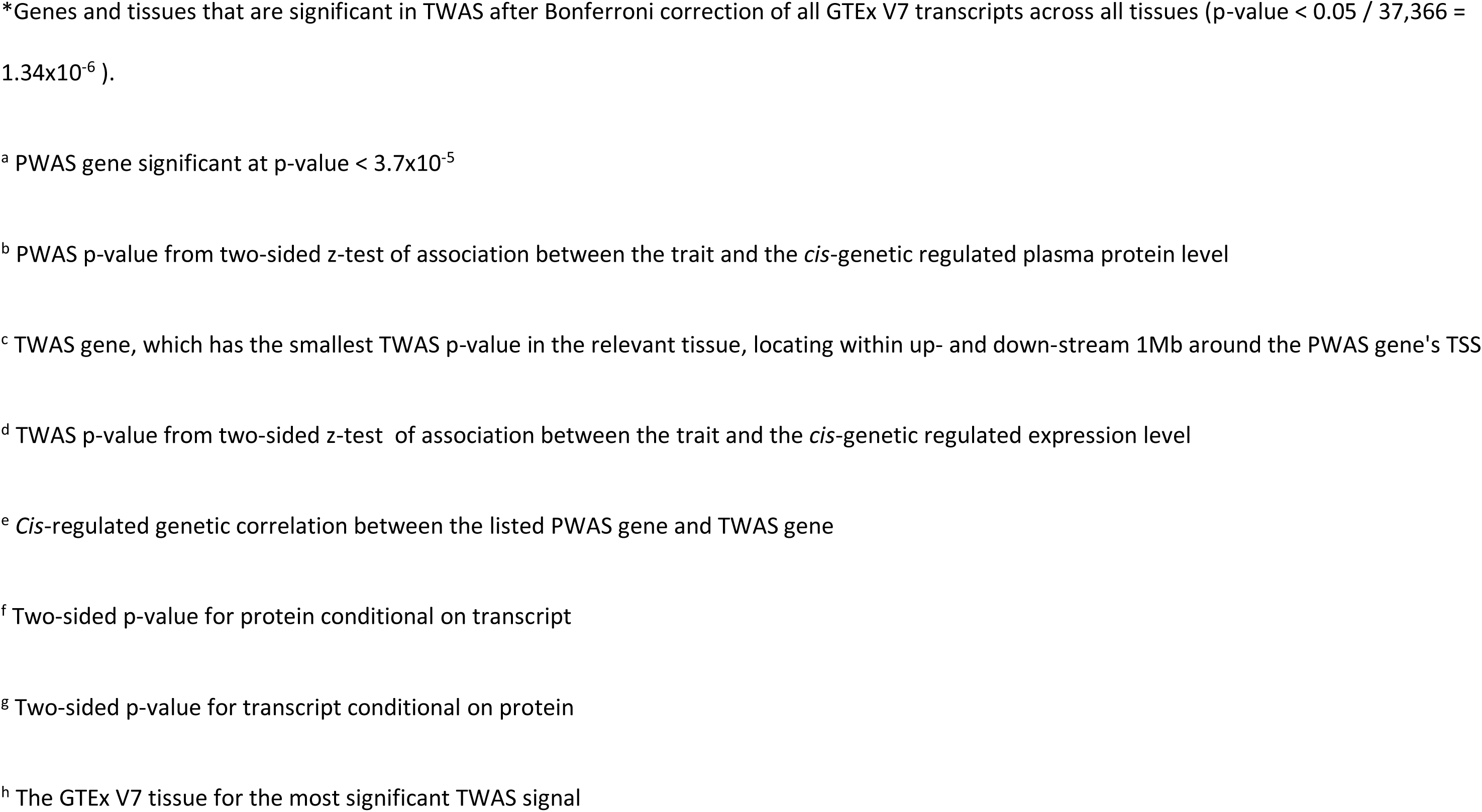
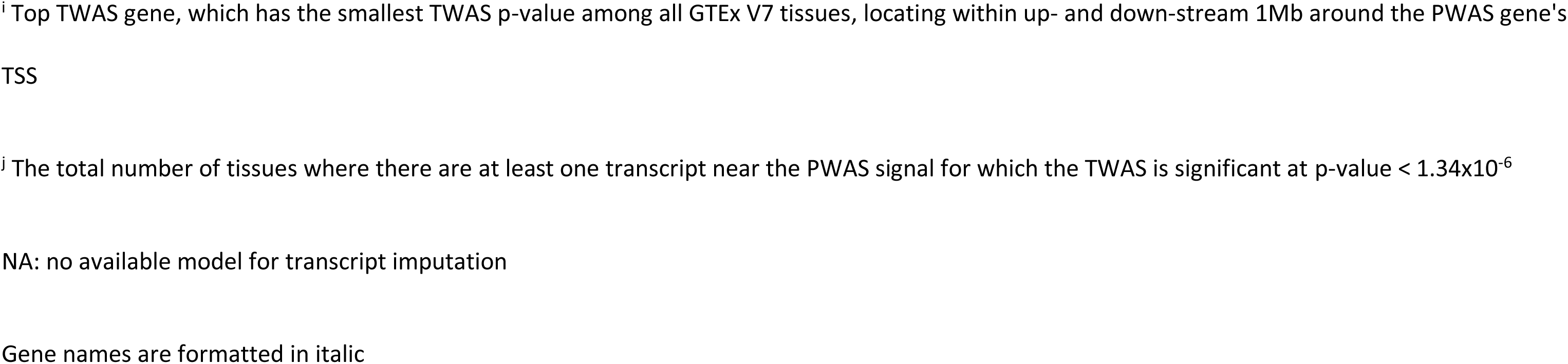
Proteome-wide association analysis of Serum Urate Level. Ten distinct loci containing significant PWAS genes (p-value < 0.05/1,348 = 3.7x10^-5^) are identified in the two-sided z-test of association. Analysis is based on summary statistics data from GWAS of serum urate level (n = 288,649) and the imputation models for plasma proteome built from the ARIC study for a total of 1,348 *cis*-heritable plasma proteins (see Supplementary Table 11). Results are also shown for the most significant genes from TWAS around +/- 500kb region of the TSS of PWAS genes for two specific trait-relevant tissues (whole blood and liver) and across all tissues. Further results from bivariate analysis of genetically imputed level of the plasma protein and that of the expression for the most significant gene from the TWAS analysis are reported in terms of conditional p-values. All TWAS analyses are performed based on models available from the GTEx V7 datasets. Results for identified top genes/tissue combinations are further validated using preliminary models available from GTEx V8 (Supplementary Table 16).

**Table 2.**
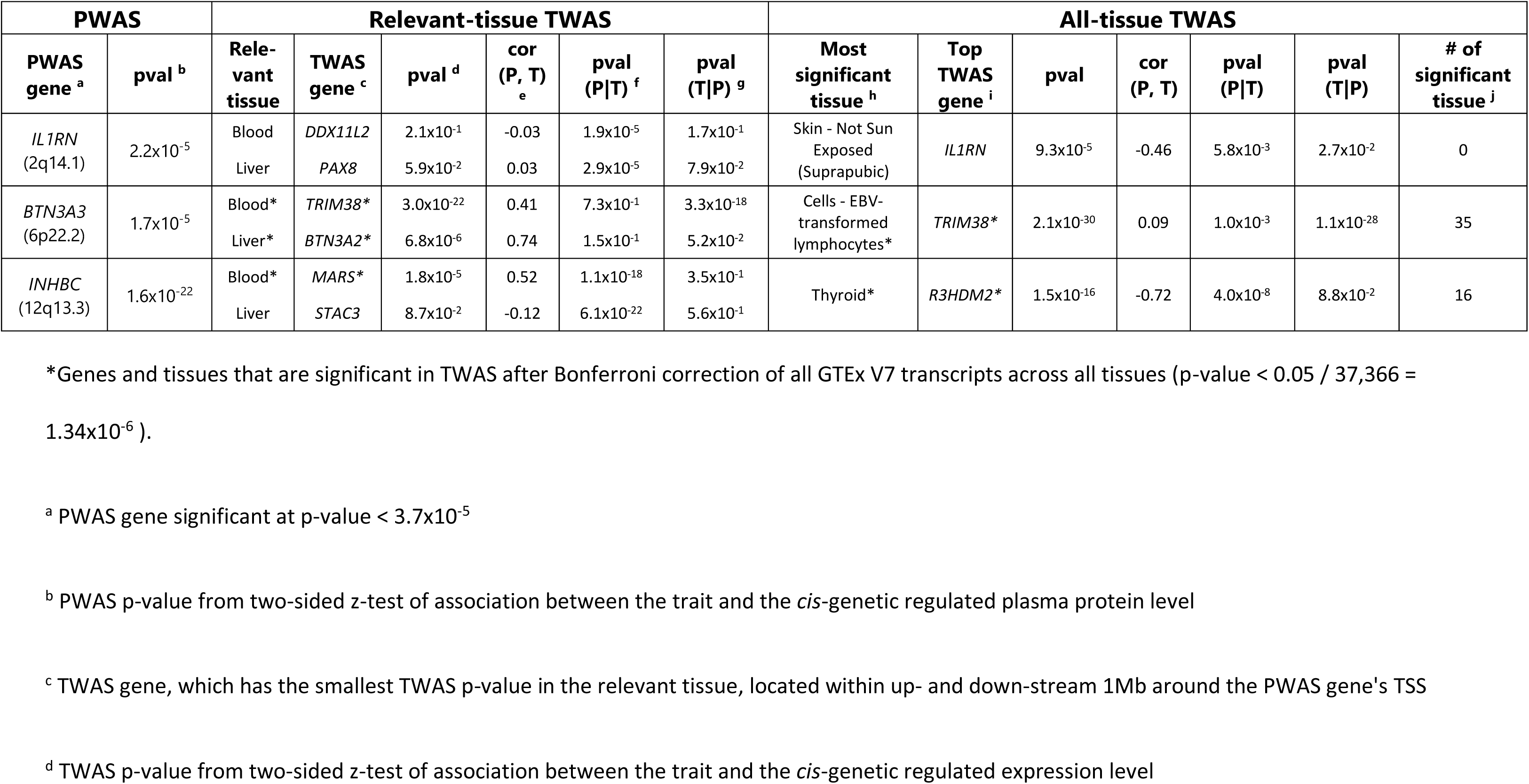
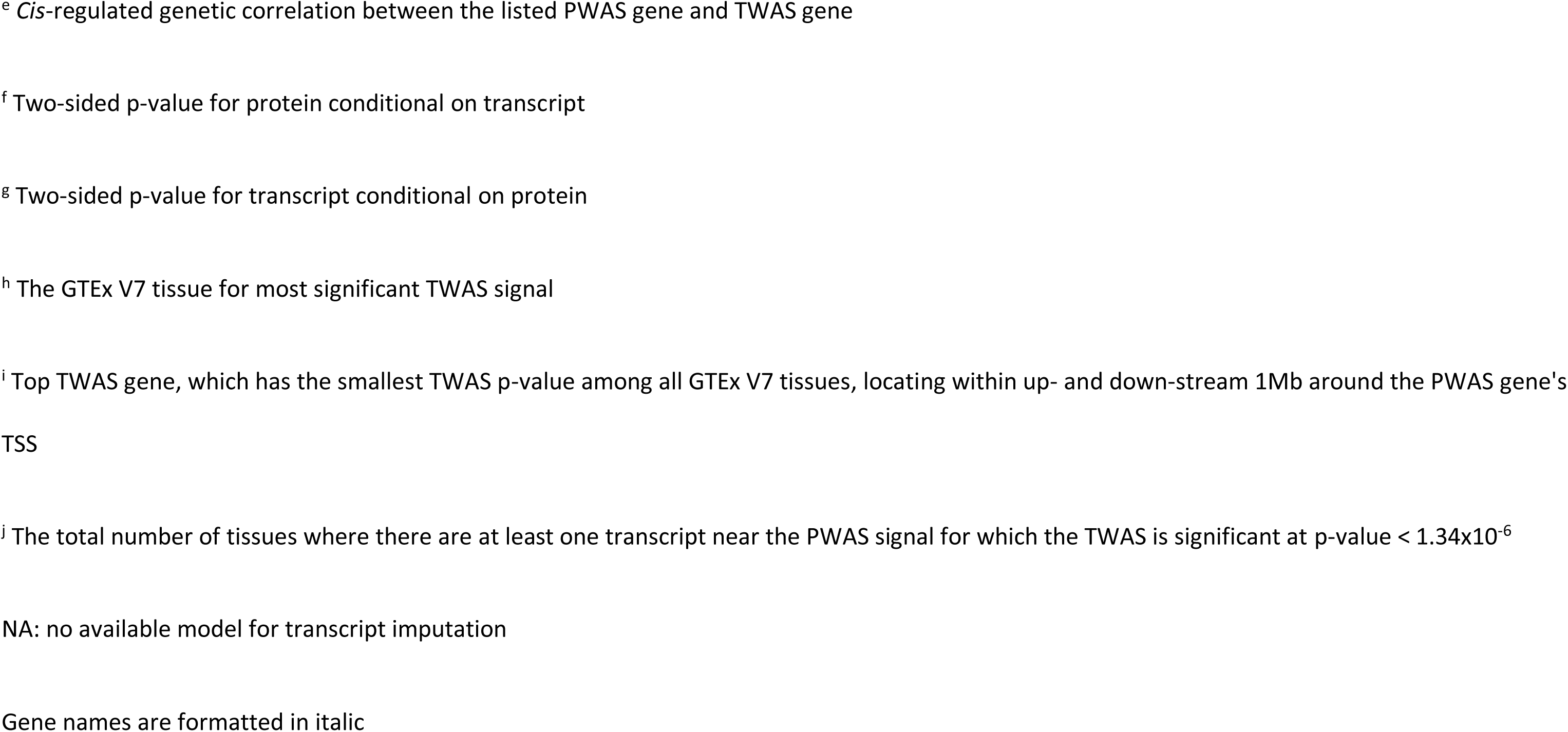
Proteome-wide association analysis of Gout. Three distinct loci containing significant PWAS genes (p-value < 0.05/1,348 = 3.7x10^-5^) are identified in the two-sided z-test of association. Analysis is based on summary-statistics data from GWAS of gout (n = 754,056) and the imputation models for plasma proteome built from the ARIC study for a total of 1,348 *cis*-heritable plasma proteins (see Supplementary Table 11). Results are also shown for the most significant genes from TWAS around +/- 500kb region of the TSS of PWAS genes for two specific trait-relevant tissues (whole blood and liver) and across all tissues. Further, results from bivariate analysis of genetically imputed level of the plasma protein and that of the expression for the most significant gene from the TWAS analysis are reported in terms of conditional p-values. All TWAS analyses are performed based on models available from the GTEx V7 datasets. Results for identified top genes/tissue combinations are further validated using preliminary models available from GTEx V8 (Supplementary Table 16).

The conditional analysis of serum urate revealed several interesting patterns (Table 1). First, there were PWAS signals that could be largely explained by nearby TWAS signals for the corresponding transcript in relevant tissues (e.g., *INHBB* in liver, and *SNUPN* in whole blood). This may be indicative of genetic loci influencing serum urate through altered gene expression and corresponding protein levels ^50^. Second, there were also PWAS signals that could be largely explained by the TWAS signal of the corresponding transcript in other tissues (e.g. *B3GAT3* in brain), but not in whole blood or liver. Such examples support the notion that the evaluation of diverse potential tissues of action is important to characterize these genetic loci. However, the TWAS effect of *B3GAT3* in brain are negative whereas the effect of its PWAS is positive. We found the opposite direction is consistent with their negative genetic correlation between plasma protein and gene-expression in those tissues. Third, for the locus around *INHBC*, the plasma PWAS signal for *INHBC* explains the most significant nearby TWAS signal *R3HDM2* in thyroid (conditional p-value of TWAS signal = 4.1x10^-1^) but not *vice* versa (conditional p-value of PWAS signal = 6.8x10^-34^). We found the patterns to be qualitatively similar when the analyses were repeated using the V8 models (Supplementary Table 16). For the significant PWAS signals, (Supplementary Tables 17.1 and 17.2) we further observed that whenever there was strong genetic correlation between plasma protein and gene expression there was also strong evidence of colocalization (e.g. *INHBB* in liver, and *B3GAT3* in brain, see Supplementary Tables 17.1 and 17.2).

Finally, the PWAS of gout revealed a finding illustrating the potential to detect potential drug targets based on the significant association with the Interleukin 1 Receptor Antagonist protein (IL1RN, p-value = 2.2x10^-5^) (Table 2). IL1RN binds to its target, the cell surface interleukin-1 receptor (IL1R1), thereby inhibiting the pro-inflammatory effect of interleukin-1 signaling. Anakinra, an anti-inflammatory drug approved to treat rheumatoid arthritis, is a recombinant, slightly modified version of the IL1RN protein examined in our study that binds to IL1R1, blocking its actions (Extended Data Figure 9). The observed association between higher levels of IL1RN protein and lower odds of gout are consistent with the beneficial effect of its synthetic analogue anakinra on other inflammatory diseases and suggest a repurposing opportunity for anakinra to treat acute gout flares. In fact, such evaluations are ongoing, with a recent randomized, double-blind, placebo-controlled trial of acute gout flares showing anakinra to be non-inferior to usual treatment ^51^. While drug delivery to plasma proteins and their cell surface receptors is easier than to other molecules such as intra-nuclear proteins, druggability of any implicated protein in our study depends on various factors such as protein structure and biological functions, and needs to be evaluated on a case-by-case basis. A systematic connection of all *cis*-heritable proteins to active drug candidates is provided as an additional resource (Supplementary Table 18).

## Discussion

We present a comprehensive analysis of *cis*-genetic regulation of the plasma proteome based on a large discovery study that include both EA and AA individuals and an additional replication study based on AA individuals. Our study almost tripled the number of genes with identified *cis*-pQTL compared to previous reports ^16, 17^ and led to understanding of unique genetic architecture of plasma proteome in the AA population. We developed models for plasma protein imputation separately for the two populations and make them publicly available to facilitate future proteome wide association studies. Using large-scale GWAS summary-statistics from two complex traits, we illustrate how PWAS can complement TWAS for the identification of causal genes, protein products and inform potential drug targets. We have created a web resource for downloading summary-statistics data and PWAS models with searchable options for exploring/viewing various results from our analyses (http://nilanjanchatterjeelab.org/pwas).

Our analysis provides several important insights into the *cis*-genetic architecture of plasma proteome. We observe that *cis*-heritability of protein levels tends to be smaller compared to those of gene expression levels in related tissues (Fig. 3a), a pattern consistent with the central dogma of DNA regulating the proteome through the transcriptome and the widespread presence of post-translational modification. Further, we observe important heterogeneity across the two populations. We found nearly 30% of the sentinel pQTLs detected in the AA population were non-existent or extremely rare in the EA population, but the converse proportion was much more modest (∼10%). We also observe that the cross-population performance of protein imputation models is better from AA to EA population than the converse (Fig. 3c). Population-specific fine-mapping analysis indicated that the size of “credible set” for many genes is substantially smaller in the AA than the EA population. Taken all together, our analysis demonstrates that similar to what has been reported earlier for complex traits ^52^, there are distinct advantages of including samples from diverse ancestries in genetic studies of molecular phenotypes.

While we increased the number of known *cis*-pQTLs , some of the patterns of associations we see have been noted earlier. For example, a prior study ^24^ has previously shown that pQTLs identified in the EA population largely replicates in non-EA Arabic and Asian population. However, besides the high degree of correlations in effect sizes for *cis*-pQTLs common across both populations, we also showed that discovery analysis in the AA population itself leads to the identification of many unique *cis*-pQTLs and further fine-mapping analysis in this population leads to better resolution for the identification of causal variants.

We demonstrate applications of protein imputation models for conducting proteome-wide association studies (PWAS) for two related complex traits, resulting in the exemplary identification of the *IL1RN* protein which indicates potential promise for drug repurposing of anakinra to treat acute gout flares. Through multivariate analysis, we further explored relationship between plasma PWAS signals and those detected at the transcriptome level through complementary TWAS approach across various tissues. We found that while TWAS signals often exist in the same region, the underlying genes for which the strongest signals are seen can differ or/and the underlying tissue may not be closely related to plasma. As plasma proteins are easier target for drug delivery, we created an additional resource connecting all *cis*-heritable proteins to active drug candidates (Supplementary Table 18). In general, we believe the most promising target genes could be where there exists both PWAS and TWAS signals with underlying evidence of genetic correlation and colocalization.

Our study has several limitations. First, while the platform we used included SOMAmers for close to 5,000 proteins or protein complexes, it does not provide coverage for the entire plasma proteome. In the future, more comprehensive protein measurements across different tissues will be needed to further pinpoint target genes and tissues of actions. Second, the power of our PWAS analysis conditional on TWAS signals may be affected by small sample size of underlying eQTL datasets. Third, in this study, we have not carried out a joint analysis of the data across the two population and thus may have incurred some loss of power for the identification of shared pQTLs. Fourth, we have not explored effects of uncommon and rare variants, as well as complex trans-associations, all of which could have significant impact in explaining heritability, but substantial discovery is likely to need even larger sample size.

In conclusion, our study, together with two other contemporary investigations ^53, 54^, provides comprehensive and cross-population insight into genetic architecture of plasma proteome. We generate several resources (http://nilanjanchatterjeelab.org/pwas) for utilizing our results to investigate the causal role of plasma proteins on complex traits and their drug repurposing potential.

## Acknowledgements

The Atherosclerosis Risk in Communities study has been funded in whole or in part with Federal funds from the National Heart, Lung, and Blood Institute, National Institutes of Health, Department of Health and Human Services, under Contract nos. (HHSN268201700001I, HHSN268201700002I, HHSN268201700003I, HHSN268201700005I, HHSN268201700004I). The authors thank the staff and participants of the ARIC study for their important contributions. SomaLogic Inc. conducted the SomaScan assays in exchange for use of ARIC data. This work was supported in part by NIH/NHLBI grant R01 HL134320. The UK BioBank data was obtained under the UK BioBank resource application 17712. Research of J.Z., D.D., and N.C. was supported R01 grant from the National Human Genome Research Institute [1 R01 HG010480-01]. B.H. was supported by Bloomberg Distinguished Professorship Endowment fund available to N.C.. The work of A.K. was funded by the Deutsche Forschungsgemeinschaft (DFG, German Research Foundation) – Project-ID 431984000 – SFB 1453. The work of P.S. was funded by the EQUIP Program for Medical Scientists, Faculty of Medicine, University of Freiburg. The work of A.T. was funded by R01 AR073178. The work of J.C. and E.B. was funded by the ARIC contract. The work of M.G. and J.C. was funded by the multiomics grant R01 DK124399. The work of B.Y. was funded by HL148218. We acknowledge Dr. Nicholas Mancuso and Dr. Alexander Gusev for providing preliminary TWAS models built with GTEx V8 data.

## Author Contribution Statement

J.Z., J.C. and N.C. conceived the project. J.Z. and D.D. carried out all data analyses with supervision from N.C.. B.H. developed online resources for data visualization and sharing, J.Z., D.D., A.K. and N.C. drafted the manuscript, and A.T., P.S., M.G. and B.Y. provided comments. All co-authors reviewed and approved the final version of the manuscript.

## Competing Interests Statement

Proteomic assays in ARIC were conducted free of charge as part of a data exchange agreement with Soma Logic. The authors declare no other competing interests.

## CKDGen Consortium

Anna Köttgen^2,3^, Adrienne Tin^2,4^, Eric Boerwinkle^6,7^, Josef Coresh^1,2,5^

Affiliations:

1. Department of Biostatistics, Johns Hopkins Bloomberg School of Public Health, Baltimore, MD, USA

2. Department of Epidemiology, Johns Hopkins Bloomberg School of Public Health, Baltimore, MD, USA

3. Institute of Genetic Epidemiology, Faculty of Medicine and Medical Center - University of Freiburg, Freiburg, Germany

4. MIND Center and Division of Nephrology, University of Mississippi Medical Center, Jackson, MS, USA

5. Department of Medicine, Johns Hopkins University School of Medicine, Baltimore, MD, US

6. Epidemiology, Human Genetics and Environmental Sciences, School of Public Health, University of Texas Health Science Center at Houston, Houston, TX, USA

7. Human Genome Sequencing Center, Baylor College of Medicine, Houston, TX, USA

## Data availability

Genome-wide summary-level statistics for all single-SNP cis-pQTL analysis, irrespective of significance level, and data required to perform PWAS, are available from http://nilanjanchatterjeelab.org/pwas. For individual-level plasma protein data, pre-existing data access policies for each of the parent cohort studies (ARIC and AASK) specify that research data requests can be submitted to each steering committee; these will be promptly reviewed for confidentiality or intellectual property restrictions and will not unreasonably be refused. Please refer to the data sharing policies of these studies. Individual level patient or protein data may further be restricted by consent, confidentiality or privacy laws/considerations. These policies apply to both clinical and proteomic data. The CKDGen Consortium makes all data reported in its original publications publicly available (https://ckdgen.imbi.uni-freiburg.de/). For European-specific gout GWAS data, additional data requests can be submitted to the CKDGen steering committee; these will be promptly reviewed for confidentiality or intellectual property restrictions and will not unreasonably be refused. GRCh38 reference genome data from Phase-3 1000 Genome Project is available from https://www.internationalgenome.org/data. Access to UK Biobank individual level data can be requested from https://www.ukbiobank.ac.uk/enable-your-research/apply-for-access. Gene expression imputation models previously built based on data from the GTEx V7 and data required to perform TWAS are available from http://gusevlab.org/projects/fusion/#reference-functional-data; models based on GTEx V8 are available on request from Dr. Nicholas Mancuso and Dr. Alexander Gusev. *cis*-eQTL summary statistics are available from https://gtexportal.org/home/. VEP was obtained from https://useast.ensembl.org/index.html. Therapeutic target database was downloaded from http://db.idrblab.net/ttd/full-data-download.

## Code availability

The codes used to perform data analysis relevant to this paper, including protein data cleaning, *cis*-pQTL mapping, building PWAS models, etc., are available from https://github.com/nchatterjeelab/PlasmaProtein. Example codes to perform PWAS using external GWAS data are available from http://nilanjanchatterjeelab.org/pwas. The majority of our statistical analysis was performed using R 3.6.1 and R 4.0.2, and R packages biomaRt 2.42.1, peer 1.0, plink2R 1.1, glmnet 4.0, ggplot2 3.3.3, gaston 1.5.6, GGally 2.0.0, ggpubr 0.4.0, readr 1.3.1, bigreadr 0.2.0, readxl 1.3.1, xlsx 0.6.3, dplyr 1.0.4, stringr 1.4.0, latex2exp 0.4.0. *Cis*-pQTL mapping was performed using QTLtools v1.2 (Binary CentOS 7.8). The publicly available summary-level statistics and analysis relevant to analyzing genotype data were performed by PLINK 2.0 and PLINK 1.9. *Cis*-heritability analysis was performed using GCTA 1.93.0 beta. Plasma protein imputation models were trained using FUSION available from https://github.com/gusevlab/fusion_twas. Downstream analysis including enrichment and colocalization was performed using VEP (version 85), TORUS (https://github.com/xqwen/torus), and coloc v3.2.1. Fine-mapping was performed using SuSIE v0.11.42 for ancestry-specific analysis, and MANTRA [1.0; Feb 2012] (available on request from Professor Andrew P. Morris) for trans-ancestry analysis.

## Methods

### Study population

Our study was conducted using individual-level data from the Atherosclerosis Risk in Communities (ARIC) study ^28^. The ARIC study is an ongoing community-based cohort study of individuals that initially enrolled 15,792 participants 1987 and 1989 from four communities across the US: Washington County, Maryland; suburbs of Minneapolis, Minnesota; Forsyth County, North Carolina; and Jackson, Mississippi. The third visit (v3) occurred in 1993-1995, when blood samples used for the measurement of the proteome were collected. A total of 9,084 participants with cleaned plasma protein data (1,871 African Americans (AA), 7,213 European Americans (EA)) after the exclusions of participants without genotype data (see below) were retained in the current study.

### Plasma protein data and genetic data

The relative concentrations of plasma proteins or protein complexes from the blood samples were measured by SomaLogic Inc. (Boulder, Colorado, US) using an aptamer (SOMAmer)-based approach ^11, 12^. Details for this approach and the SomaLogic normalization pipeline can be found in a technical white paper on the manufacturer’s website, http://somalogic.com/wp-content/uploads/2017/06/SSM-002-Technical-White-Paper_010916_LSM1.pdf, and https://somalogic.com/wp-content/uploads/2017/06/SSM-071-Rev-0-Technical-Note-SOMAscan-Data-Standardization.pdf. Of the 4,877 SOMAmers measuring 4,697 unique proteins or protein complexes, we excluded 43 SOMAmers that mapped to multiple gene targets, 9 SOMAmers whose target proteins’ encoding genes do not have position record in the biomaRt database ^55^, and 8 SOMAmers without any SNPs in *cis* region. By restricting analysis to plasma proteins or protein complexes encoded by autosomal genes, we further excluded 158 genes on the X chromosome, and 2 genes on the Y chromosome. In total, 4,657 SOMAmers measuring 4,483 unique proteins or protein complexes encoded by 4,435 autosomal genes passed quality control, and were retained in the current study.

Genotyping of ARIC samples was performed on the Affymetrix 6.0 DNA microarray and imputed to the TOPMed reference panel (Freeze 5b) ^56, 57^. The SNPs with imputation quality R^2^ < 0.8, call rates <90%, Hardy-Weinberg equilibrium p-values < 10^-6^, or minor allele frequencies <1% were excluded. Genetic principal components show that the two self-reported ancestry, European Americans (EA) and African Americans (AA) are well distinguished in terms of genetic ancestry (**Extended Data Figure 10**) ^58^.

### Plasma protein data processing

Additional variation in high-throughput gene expression data which is not due to genetic variants has been found to impact the power of eQTL discoveries ^8, 9^. The fluctuations of internal environment, experimental deviations, and batch effects can all have large influence on high throughput measurements ^32^. To study whether this type of variance exists in our high-throughput plasma protein data measured by the SOMAmers, we performed analysis of variance (ANOVA) test for non-genetic factors to the first 10 principal components (PCs) of log-transformed relative abundance of SOMAmers. Non-genetic factors include common covariates (age, sex, and study sites at v3), as well as batch effects (plate run date, scanner ID, plate position, and subarray). (**Supplementary Table 19**).

To account for those non-genetic variances, which may obscure genetic association signals, we used the Probabilistic Estimation of Expression Residuals (PEER) method to estimate a set of latent covariates, and put them linearly in the model ^33^. The number of PEER factors for each ancestry was selected to maximize the number of significant SOMAmers, i.e. SOMAmers with a significant *cis*-pQTL near the putative protein’s gene.

The log-transformed relative abundance of SOMAmers were adjusted in a linear regression model including PEER factors and the covariates sex, age, study site, and 10 genetic principal components (PCs). The residuals from this linear regression were then rank-inverse normalized to avoid the influence of extreme values, and were used as the corrected-protein quantification in the analysis. By analyzing up to 200 PEER factors in increments of 10, the maximum of number of significant SOMAmers were achieved at 90 and 80 PEER factors for EA and AA populations, respectively (**Fig. 1a**). Thus, the corrected-protein quantifications adjusted for 90 and 80 PEER factors were used as phenotypes in the analysis of the EA and AA populations, respectively.

### Significant SOMAmers discovery

Significant SOMAmer is defined as SOMAmer with a significant *cis*-pQTL near the putative protein’s gene. For all primary analyses, we defined the mapping window as 500-kb upstream and downstream of the target protein-coding genes’ transcription start site (TSS). In a secondary analysis, we found that *cis*-heritability of SNPs within +/- 500Kb and +/- 1Mb of the TSS to be quite similar, indicating that vast majority of *cis*-pQTLs for the larger region to be concentrated within +/- 500Kb window (**Supplementary Table 20**). Gene position of GRCh38 reference genome was obtained from Ensembl BioMart database ^55^. Common linear regression procedures for association tests using the Bonferroni correction to p-values usually proves to be overly stringent and results in many false negatives ^38^. To overcome this issue, adaptive permutation approach implemented in QTLtools were applied ^37^. We used one hundred permutations to empirically characterize the null distribution of the strongest signal which is fitted by a Beta distribution. The p-values of association adjusted for the number of variants tested in *cis* given by the fitted beta distribution were used to derive SOMAmer-level (gene-level) nominal p-values. By controlling the false discovery rate (FDR) threshold < 5%, significant SOMAmers were identified.

### Comparison with previous identified *cis*-pQTL

A list of existing pQTL studies were summarized by Karsten Suhre (http://www.metabolomix.com/a-table-of-all-published-gwas-with-proteomics/) ^24^. We focus on two recent European-ancestry pQTL studies with large sample size and proteins assayed by SOMAscan. The first was performed in the INTERVAL study with UK blood donors ^15^. The other was performed in the AGES-RS cohort ^16^. To make fair comparison, we compared identified *cis*-pQTLs across the two analyses using the same standard -- sentinel *cis*-associations (+/-500Kb) for common SNPs (MAF>0.01) and Bonferroni corrected genome-wide threshold for significance (p-value < 1.5x10^−11^ in INTERVAL, and 1.92×10^-10^ in AGES-RS). Using these criteria, the two previous studies identified a total of 508 unique significant SOMAmers (304 and 422 respectively) and we identified 1,465 significant SOMAmers. We then tested replication of their sentinel SNPs in our ARIC EA sample (Bonferroni corrected p-value < 0.05/726 = 6.89x10^-5^, where 726 = 218x2 + 204 + 86. There were 218 SOMAmers discovered in both studies, 204 discovered only in AGES-RS and 86 discovered only in INTERVAL). If a significant SOMAmer’s sentinel SNPs was not available in ARIC, we used their LD proxies and the *r^2^* was calculated from the 1000Genome European individuals.

### Replication of *cis*-pQTL identified in AA

We replicated *cis-*pQTLs discovered in the ARIC AA in the African American Study of Kidney Disease and Hypertension (AASK), a clinical trial of alternate blood pressure lowering regimen and goals ^35^. Enrollment occurred from 1995 to 1998, with the original trial population consisting of 1094 African American participants with chronic kidney disease. Blood samples used for the measurement of the proteome were collected at baseline. A total of 467 participants with serum protein data and genotype data were retained in the current study. Proteomic profiling was performed using the SomaScan technology using the V4.1 platform. Genotyping was conducted using the Infinium Muti-Ethnic Global BeadChip array (Illumina, GenomeStudio) and imputed to the TOPMed reference panel (Freeze 5 on GRCh38).

### Independent *cis*-pQTL mapping

It is likely that the significant SOMAmers have multiple proximal *cis*-SNPs which have independent effects. To identify independent signals for them, we performed independent *cis*-pQTL mapping using the conditional pass implemented in QTLtools ^37^. The algorithm first uses permutations to derive SOMAmer-level (gene-level) nominal p-values (as described in **Significant SOMAmers discovery**), then it uses a forward-backward stepwise regression to select the conditional independent signals using this significance threshold. In this process, it automatically learns the number of independent signals per SOMAmer using forward selection, and then determines the best candidate SNP per signal using backward selection controlling for the remaining signals. If no SNP is significant at the previous nominal p-value threshold, the candidate signal will be dropped; otherwise, the SNP with smallest backward-p-value will be chosen as the lead SNP for this candidate signal. In some cases, the same SNP during the backward selection can explain multiple independent signals that were detected during the forward selection. In the reporting our results (**Supplementary Table 6.1 and 6.2**), we show the rank of all the SNPs selected by the forward selection step that is explained by a given lead SNP selected during the final backward selection step.

To account for power for detection in **Fig. 1c**, we adjusted the SNP effect sizes by assigning a weight of the inverse of statistical power. The statistical power can be derived as following. The SNP effect is chi-square distributed with one degree of freedom (df). It is a central chi-square distribution under the null, and a non-central chi-square distribution under the alternative hypothesis. The non-centrality parameter (NCP), 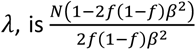 where *N* is the number of samples in study, *f* is the MAF of the SNP, and *β* is the SNP effect ^59, 60^. The significance threshold for the test statistic under the central chi-square distribution of df 1 and the SOMAmer’s nominal p-value cut-off, *p*_0_, is *t*_0_ = F^−1^(1−*p*_0_, 1), where *F*(·,1) is the cumulative distribution function (CDF) of a central chi-square distribution of df 1. The statistical power can be computed by *Pr*(*T* > *t*_0_|*H_a_*) = 1 – *G*(*t*_0_,*λ*,1), where *T* is the test statistics and *G*(·, λ, 1) is the CDF of the non-central chi-square distribution with NCP of *λ* and df 1. The weight assigned to SNP effect is (1 – *G*(*t*_0_,λ,1))^−1^.

### Investigation of epitope-binding effects

SOMAscan assay relies on aptamer binding which may be influenced by the change of protein structure. Protein altering variants (PAV) may result in *cis*-pQTLs by altering binding affinity, instead of protein abundance. Following a procedure recommend earlier ^15^, we cataloged all *cis*-pQTLs that were not in LD (*r^2^*<0.1) with any PAV in the *cis* region or those in LD (0.1≤*r^2^*≤0.9) but remain significant in a conditional analysis after adjusting for PAVs. We annotated variants with variant effect predictor (VEP) ^61^, Loss-Of-Function Transcript Effect Estimator (LOFTEE) ^62^ and Ensembl Regulatory Build ^63^. Variants were considered to be PAV if annotated as coding sequence, frameshift, in-frame deletion, in-frame insertion, missense, splice acceptor, splice donor, splice region, start lost, stop gained, or stop lost variants. LD-pruned (*r^2^*>0.9) PAVs were included as covariates for association testing.

### *Cis*-eQTL overlap

We cross referenced the identified *cis*-pQTLs against *cis*-eQTLs identified in the overall analysis of GTEx (V8) data across different tissues. For each SOMAmer, we first extracted the sentinel *cis*-pQTLs, meaning the variants having most significant association along with all the variants in high LD (*r^2^* > 0.8). Using this list of variants across 2,004 SOMAmers which had at least one *cis*-pQTL in EA, we calculated the percentage overlap with the set of significant *cis*-eQTLs (at FDR<5%, as defined by GTEx consortium) for the same gene identified in each tissue of GTEx V8 ^9^. Since the GTEx cohort is primarily of European ancestry, we restricted this analysis to EA only.

### Colocalization

Colocalization analysis was performed to investigate whether the same variants were likely to be causal for variation in protein levels and gene expression levels. We used publicly available overall *cis*-eQTL summary statistics from GTEx consortium (V8). For testing whether *cis*-eQTL and *cis*-pQTL associations for the same gene colocalize, we used coloc package in R with the default setting ^64^. Evidence for colocalization was assessed using the posterior probability (PP) for the hypothesis that there is an association for both protein levels and gene expression levels, and they are driven by the same causal variant (PP.H4). Since we tested across a large number of tissues, we chose a stringent cut-off of 0.8 and significant SOMAmers with PP.H4 > 0.8 were identified as likely to have a shared causal variant for the *cis*-eQTL and *cis*-pQTL associations. As before, we restricted our analysis to the 2,004 significant SOMAmers identified in EA.

### Function annotations enrichment

We performed an enrichment analysis of the *cis*-pQTLs for known regulatory elements in the genome to identify the broad functions of the *cis*-pQTLs. The functional annotations were curated from variant effect predictor (VEP) ^61^, Loss-Of-Function Transcript Effect Estimator (LOFTEE) ^62^ and Ensembl Regulatory Build as was reported in the recent GTEx analysis. For each SOMAmer, we used sentinel *cis*-pQTLs, meaning the variants having the most significant association and variants in high LD (*r^2^* > 0.8) for evaluating functional enrichment. With these annotations, we used TORUS ^65^ to perform functional enrichment for each functional category. TORUS uses a hierarchical Bayesian approach to integrate genomic annotations in QTL mapping. In particular, it uses a logistic prior to model the enrichment of a genomic annotation and employs an EM-algorithm based approach to perform inference on the enrichment parameters. Further it outputs the 95% confidence intervals of the log enrichment parameters from which the p-value can be calculated under asymptotic normality assumptions. The details have been outlined in Wen 2016 ^65^. To remove effect of potential epitope binding effects associated with the PAVs, we also investigated functional enrichment among sentinel *cis*-pQTLs (and variants in high LD) that showed significant effects independent of the PAVs (See previous section for details).

### Fine-mapping analysis

To identify the set of possibly causal variants regulating plasma protein levels we performed fine-mapping ^66^ using the *cis*-variants for each of the 1,447 SOMAmers that had at least one *cis*-pQTL in both populations using SuSiE ^39^. SuSiE uses a single effect regression model, with normal and multinomial prior distributions for effect sizes and inclusion probabilities and subsequently employs a variational approximation to compute the posterior probabilities. Under the default settings, SuSiE assumes that each genetic variant has the same probability of inclusion in the credible set (see Wang et al. ^39^ for details). For a given SOMAmer and corresponding variants in the *cis*-regulatory region, SuSiE outputs a number of single effect components or credible sets that have 95% probability to contain a variant with non-zero causal effect. We set the maximum number of such singlet effect components to be 10, meaning broadly we allow for the possibility that a SOMAmer can be regulated by 10 causal variants at best. Further, SuSiE also outputs the posterior inclusion probability for each variant. This corresponds to the probability of the variant to be included in one of the credible sets.

To perform trans-ancestry meta-analysis, we used MANTRA ^40^ which is based on a computationally intensive Bayesian partition accounting for the shared similarity in closely related populations assuming the same underlying allelic effect. It models the effect heterogeneity among distant populations by clustering according to the shared ancestry and allelic effects. Under the default setting the prior density for the effect sizes is given by a normal distribution and the prior for the number of clusters is given by a mixture geometric distribution. MANTRA outputs the Bayes factor for association of a variant across ancestries. Using this, we constructed the posterior probability ^67^ of the k^th^ variant (*π_k_*) as:

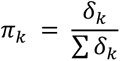

where *δ_k_* is the Bayes factor for association of the kth variant obtained using trans-ancestry meta-analysis in MANTRA and the sum in the denominator is across all the variants in the *cis*-region. We performed MANTRA using the variants common to EA and AA and subsequently calculated the posterior probabilities.

### *Cis*-SNP heritability estimation

*Cis*-SNP heritability (*cis*-h^2^) of SOMAmers were estimated using the REML algorithm implemented in GCTA ^46^. Genotypes of SNPs in a *cis*-window around the encoding gene of the corresponding target protein of a SOMAmer were used to estimate genetic relatedness matrix (GRM). Corrected-protein quantifications and the estimated GRM were input to the GCTA to estimate *cis*-h^2^ using the REML algorithm (option --reml --reml-no-constrain). A maximum number of 100 iterations was set to determine the convergence of the estimation algorithm. The nonzero *cis*-heritability was tested using a likelihood-ratio test for the first genetic variance component (option --reml-lrt 1) with significance level of 0.01. Plasma protein SOMAmers with negative estimate *cis*-h^2^ estimates were excluded. *Cis* window size of +/- 500Kb and 1Mb were examined, and there were no significant differences between the heritability estimations (**Supplementary Table 20**). Therefore, throughout the paper, we defined +/- 500Kb window size which is same as those used for TWAS models we used.

### Imputation models trained jointly with *cis*-SNPs

Using the TWAS / FUSION ^27^ (http://gusevlab.org/projects/fusion/), we built imputation models for 1,394 (AA) and 1,350 (EA) SOMAmers with significant non-zero *cis*-h^2^. Imputation model for a SOMAmer was trained jointly by elastic net using *cis*-SNPs in +/-500Kb around the TSS of the encoding gene of the target protein. The tuning parameters were selected based on 5-fold cross-validation, and the final elastic net model was re-fitted using all data and the selected tuning parameters. The coefficients for SNPs were all zero in the re-fitted elastic net models for nine SOMAmers in AA and two SOMAmers in EA, respectively. So these proteins were excluded in the following analysis (see **Supplementary Table 11**). The performance of models was evaluated by adjusted prediction accuracy which was defined as the 5-fold cross-validated R^2^ between predicted and true values standardized by *cis*-h^2^. The imputation models built only with the sentinel *cis*-pQTL was used as a baseline comparison.

### Trans-ancestry prediction capacity

To study the trans-ancestry prediction performance, we applied the genetic imputation models to the genotypes of individuals from their opposite races in ARIC. The cross-ancestry prediction performance is evaluated by the R^2^ between predicted and true values standardized by *cis*-h^2^.

### *Cis*-regulated genetic correlation between plasma proteome and transcriptome across a variety of tissues

To study the *cis*-regulated genetic correlation between plasma protein and expression levels for underlying genes across a variety of tissues, we computed the Pearson’s correlation coefficients between genotypically-imputed plasma proteins and genotypically-imputed gene expressions for the same gene for individuals from Phase-3 1000 Genome Project (1000Genome) ^34^ by applying weights of their imputation models to the genotype data. For primary analyses, we used established gene expression imputation models available based on GTex V7 dataset across different tissues (http://gusevlab.org/projects/fusion/#reference-functional-data (see **Supplementary Table 13** for the full list, **Supplementary Table 14** for their prediction accuracies). Here we only studied for genes significant *cis*-heritable (p-value of *cis*-h^2^ from GCTA < 0.01) for both gene expression levels and plasma protein levels (**Supplementary Tables 15.1 and 15.2**). Since the gene expression imputation models were derived using participants predominantly from European ancestry from GTEx V7, the plasma protein imputation models here were restricted to EA-derived only. If multiple transcripts or SOMAmers were measured for the same gene, the sum of their imputed levels was used to represent “the total level of the gene” in terms of gene expression or plasma protein level. We also obtained preliminary gene-expression imputation models trained based on GTEx V8 dataset (obtained based personal communication with Gusev lab) and used them to conduct several secondary/validation analyses for comparison of results with V7.

### Proteome-wide association studies (PWAS)

As an analog of TWAS, weights in the imputation models of SOMAmers can be applied to summary level data using the test statistics derived in TWAS / FUSION (http://gusevlab.org/projects/fusion/). The mathematical derivation can be found in the original paper ^27^. The type 1 error of PWAS is well-controlled in simulation using null phenotypes simulated from UK Biobank using 337,484 unrelated European ancestry individuals ^68^. As mentioned before, the coefficients for SNPs were all zero in the re-fitted elastic net models for nine SOMAmers in AA and two SOMAmers in EA, respectively. After excluding them, 1,385 (AA) and 1,348 (EA) imputation models were available in PWAS. The significance level for PWAS loci identification is adjusted by of the total number of imputation models for significant *cis*-heritable plasma proteins or protein complexes (p-value < 0.05/1,348=3.7x10^-5^ in EA which was used in our PWAS of serum urate and gout). As discussed in a recent TWAS paper ^47^, multiple SOMAmers, whose encoding genes of their target proteins or protein complexes locate closely in a locus, were sometimes identified at the same time. To identify distinct loci, a 1Mb region (+/- 500Kb of TSS) was defined around each encoding gene of the target protein of significant SOMAmers, and overlapping regions were merged. The sentinel association in each locus was selected to be the most significant PWAS gene for this region (**Supplementary Tables 21.1 and 21.2**).

We obtained standardized estimate for the causal effect 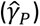 and standard error (*se*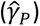), and thereby confidence intervals, of the underlying proteins on the complex traits (*Y*) by slightly extending S-PrediXcan ^69^. We derived these as

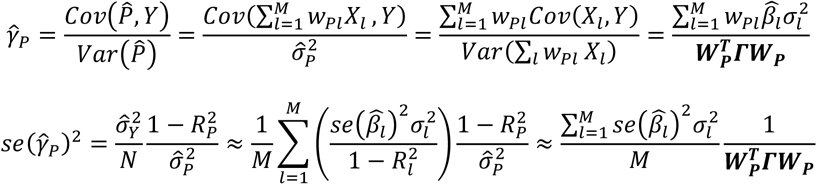

where 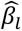 is SNP *l*’s summary statistics for the complex trait, *w_Pl_* is SNP *l*’s weight in the imputation model for protein *P,* 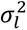 is the variance of SNP *l* which can be computed from allele frequency, and *Γ* is the LD (correlation) matrix for all *M* SNPs in the imputation model. We used the same formulae to derive corresponding causal effects, standard errors and confidence intervals for results from TWAS analyses.

### Druggability of PWAS genes

PWAS genes were annotated based on the therapeutic target database ^70^. Only drugs that were actively pursued were retained in the database and discontinued, terminated or withdrawn drugs were excluded. Additionally, druggability tiers from Finan et al. ^71^ were mapped via gene symbols (**Supplementary Table 18**).

### Bivariate conditional analysis for PWAS and TWAS

For each significant PWAS loci, we searched all TWAS genes nearby (+/-500Kb around) whose TSS locate within 500Kb of the TSS of its sentinel PWAS gene, and selected the one with the smallest TWAS p-value. The position of genes in TWAS (based on GTEx V7 based on genome build GRCh37) and PWAS (based on genome build GRCh38) were matched using the UCSC genome browser webtool (https://genome.ucsc.edu/cgi-bin/hgLiftOver) ^72^.

We first performed the nearby TWAS in two trait-relevant tissues, whole blood and liver, for serum urate and gout. Note that kidney is also a trait-relevant tissue, but there is no imputation model trained with GTEx V7 data available on TWAS / FUSION for kidney. The significance of the nearby TWAS gene was determined by significance level after Bonferroni Correction 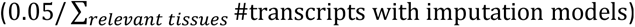.

Using z-scores (*z_p_* for PWAS gene and *z_T_* for TWAS gene) and the *cis*-regulated genetic correlation (*ρ*) of each PWAS gene and the most significant TWAS gene nearby, we performed conditional analysis ^73^ to study the potential underlying mechanism of gene expressions in tissue or proteins in plasma. The *cis*-regulated genetic correlation was computed from the Pearson’s correlation coefficients between genotypically-imputed plasma proteins and genotypically-imputed gene expressions for individuals from 1000Genome by applying weights of their imputation models to the genotype data. The least-squares estimate of the PWAS z-score conditional on TWAS z-score is

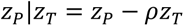

and its variance is

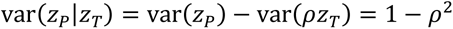

So the conditional z-score of the PWAS gene is

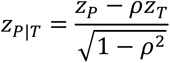

Similarly, the conditional z-score of the nearby TWAS gene is

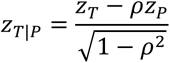

We then performed the same procedure for all nearby TWAS genes in *all* GTEx V7 tissues. Using Bonferroni Correction for the total number of transcripts with imputation models 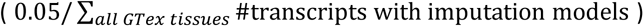, we identified the tissues which have at least one significant TWAS gene in the PWAS significant loci. The most significant TWAS gene in this region and its corresponding tissue were recorded, and then used to perform conditional analysis (**Supplementary Tables 22.1 and 22.2**). We further validated the top gene-tissue combination identified through TWAS models in V7 using preliminary models that were available to us based on V8.

## Notes

http://nilanjanchatterjeelab.org/pwas/

### Summary of Updates

Formatting and cleaning tables and figures.

