## Supplementary figures for "Plasma proteome analyses in individuals of European and African ancestry identify *cis*-pQTLs and models for proteome-wide association studies"

### Inventory of Supporting Information

| **Figure #** | **Figure title**<br>One sentence only | **Filename**<br>This should be the name the file is saved as when it is uploaded to our system. Please include the file extension. i.e.: *Smith_ED_Fig1.jpg* | **Figure Legend**<br>If you are citing a reference for the first time in these legends, please include all new references in the main text Methods References section, and carry on the numbering from the main References section of the paper. If your paper does not have a Methods section, include all new references at the end of the main Reference list. |
|---|---|---|---|
| **Extended Data Fig. 1** | *Cis*-pQTLs' effect sizes across two populations | ExtendedDataFigure1.eps | Effect sizes for common (MAF> 0.01) sentinel *cis*-pQTLs across EA and AA populations. Each dot represents a common sentinel SNP detected through either the EA (left panel) or the AA population (right panel). x-axis shows the effect size in the population through which the *cis*-pQTL is identified, and y-axis shows effect size in the other population. Minor allele frequencies (MAF) are checked for some outliers corresponding to large difference in allele frequency across populations (marked with orange). Red line is diagonal. |
| **Extended Data Fig. 2** | Overlap and colocalization of *cis*-pQTLs and *cis*-eQTLs | ExtendedDataFigure2.eps | (a) Proportion of sentinel *cis*-pQTLs in EA (including their LD-proxies; SNPs with LD > 0.8) that are identified as *cis*-eQTLs across 49 different tissues in GTEx V8. Results are ordered by the size of overlap. (b) proportion of SOMAmers showing high colocalization probability (PP.H4 > 0.8) of underlying *cis*-pQTLS and *cis*-eQTLs in the same gene across tissues in GTEx (V8). Results are ordered by the size of overlap reported in (a) for ease of comparison. |
| **Extended Data Fig. 3** | *Cis*-pQTLs tended to be significant *cis*-eQTLs across multiple tissues | ExtendedDataFigure3.eps | Distribution of number of tissues with significant *cis*-eQTL effects in GTEx V8 for the sentinel *cis*-pQTLs (and SNPs in high LD) (blue) compared to that of *cis*-eQTLs in GTEx V8 irrespective of their *cis*-pQTL status (red). Sentinel *cis*-pQTLs are restricted to those which show *cis*-eQTL effect in |

| | | | |
|---|---|---|---|
| | | | at least one tissue. *cis*-eQTL effects are evaluated for the same underlying genes for which significant *cis*-pQTLs are detected. |
| **Extended Data Fig. 4** | Functional enrichment | ExtendedDataFigure4.eps | Functional enrichment of all sentinel *cis*-pQTLs and SNPs in high LD with them ($r^2 > 0.8$) for EA (a) and AA (b). Functional enrichment of sentinel *cis*-pQTLs which have effects independent of protein altering variants are shown for EA (c) and AA (d). The red dots denote the estimated log2-enrichment statistic, and the black lines represent the corresponding 95% confidence intervals using TORUS (See Methods for details). Sample sizes for EA and AA population are n=7,213 and 1,871, respectively. |
| **Extended Data Fig. 5** | *Cis*-heritability comparison between gene expression and plasma protein levels | ExtendedDataFigure5.eps | Comparison of *cis*- heritability (*cis*-$h^2$) estimates of plasma protein (P) and gene expression (T) for a common set of overlapping genes. For each population, the overlap is defined by the set of genes that have significant *cis*-$h^2$ for both plasma protein and gene expression in the given tissue (liver and whole blood) in GTEx (a) V7 and (b) V8. Sample sizes for EA and AA populations are n=7,213 and 1,871, respectively. In boxplots, the boxes are drawn from first and third quartiles, with the median at the center, and the whiskers extending to 1.5 times the interquartile range from the box boundaries. Figures are truncated in the y-axis at *cis*-$h^2$=0 and 0.5 for better display. |
| **Extended Data Fig. 6** | Correlation between imputed gene expression and measured plasma protein levels in ARIC EA samples | ExtendedDataFigure6.eps | Measured plasma protein levels are pre-processed by inverse-rank normalization and adjusted for covariates and 90 PEER factors. Gene expression imputation models for TWAS analyses across all tissues are built based on GTEx V7 datasets (see Supplementary Table 13 for available sample sizes). The imputation models for plasma proteins are built based n=7,213 EA individuals in the ARIC study. In boxplots, the boxes are drawn from first and third quartiles, with the median at the center, and the whiskers extending to 1.5 times the interquartile range from the box |

| | | | |
|---|---|---|---|
| | | | boundaries. Figure is truncated in the y-axis at correlation= -0.15 and 0.45 for better display. |
| **Extended Data Fig. 7** | Control of type-1 error of PWAS | ExtendedDataFigure7.jpg | Quantile-quantile plot (red diagonal line) of p-values are shown for a continuous phenotype that is simulated under the null hypothesis of no genetic association for unrelated European ancestry individuals in the UK Biobank study (n=337,484). Results are based on two-sided z-tests of association between the *cis*-genetic regulated plasma protein level and the simulated null trait. The diagonal line represents expected p-values under the null hypothesis of no genetic association and the 95% confidence band, which is calculated based on standard errors of order statistics under normal approximation, represents regions of uncertainty in the q-q plot under the null hypothesis of no association. |
| **Extended Data Fig. 8** | PWAS of serum urate level and gout | ExtendedDataFigure8.jpg | Quantile-quantile plots of PWAS p-values obtained from two-sided z-tests of association between the *cis*-genetic regulated plasma protein levels and the trait of interest, serum urate level (n=288,649) and gout (n=754,056). The diagonal lines represent expected p-values under the null hypothesis of no genetic association and the 95% confidence bands, which is calculated based on standard errors of order statistics under normal approximation, represent regions of uncertainty in the q-q plot under the null hypothesis of no association. |
| **Extended Data Fig. 9** | PWAS identify repurposing opportunity for anakinra to treat gout | ExtendedDataFigure9.eps | Blue particle is interleukin-1 (IL-1) which produces pro-inflammatory effect of interleukin-1 signaling. Green particle is interleukin-1 receptor antagonist protein (IL1RN) which competes for binding but does not lead to a signal. Red particle is anakinra which has same shape as IL1RN and can also bind to the IL1R1 without eliciting a signal. Plot was created with BioRender.com. |

| | | | |
|---|---|---|---|
| **Extended Data Fig. 10** | Top five genetic principal components (PC) of ARIC data | ExtendedDataFigure10.jpg | Genetic PCs represent the major population structure in the aggregated sample of EA (blue) and AA (green) populations, colored by self-reported ancestry. |

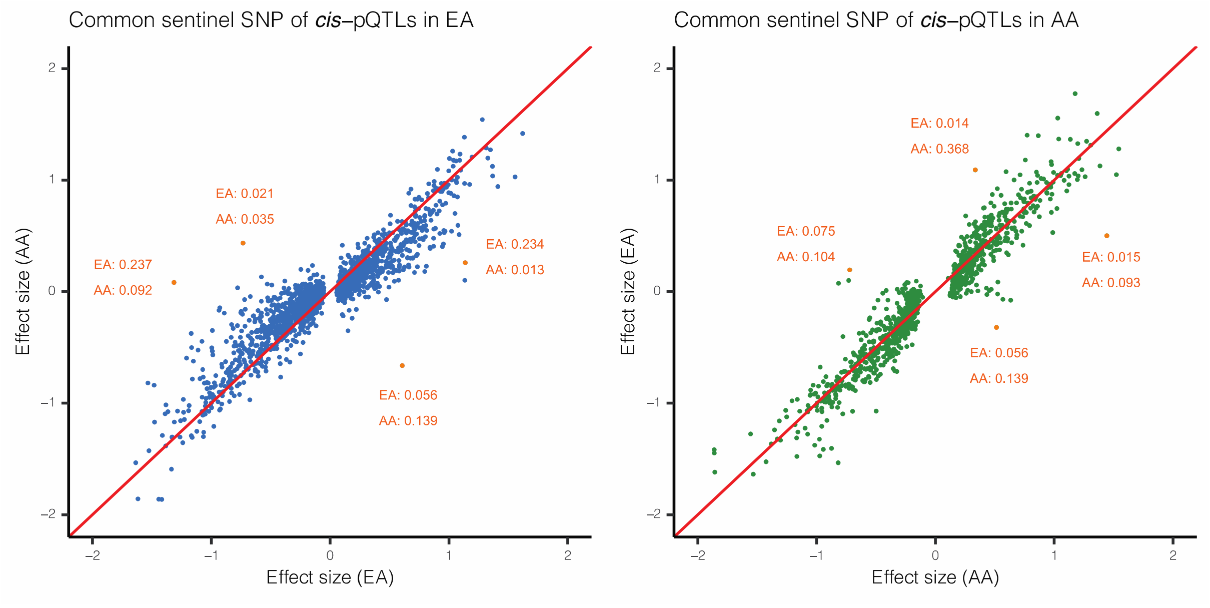


Extended Data Fig. 1

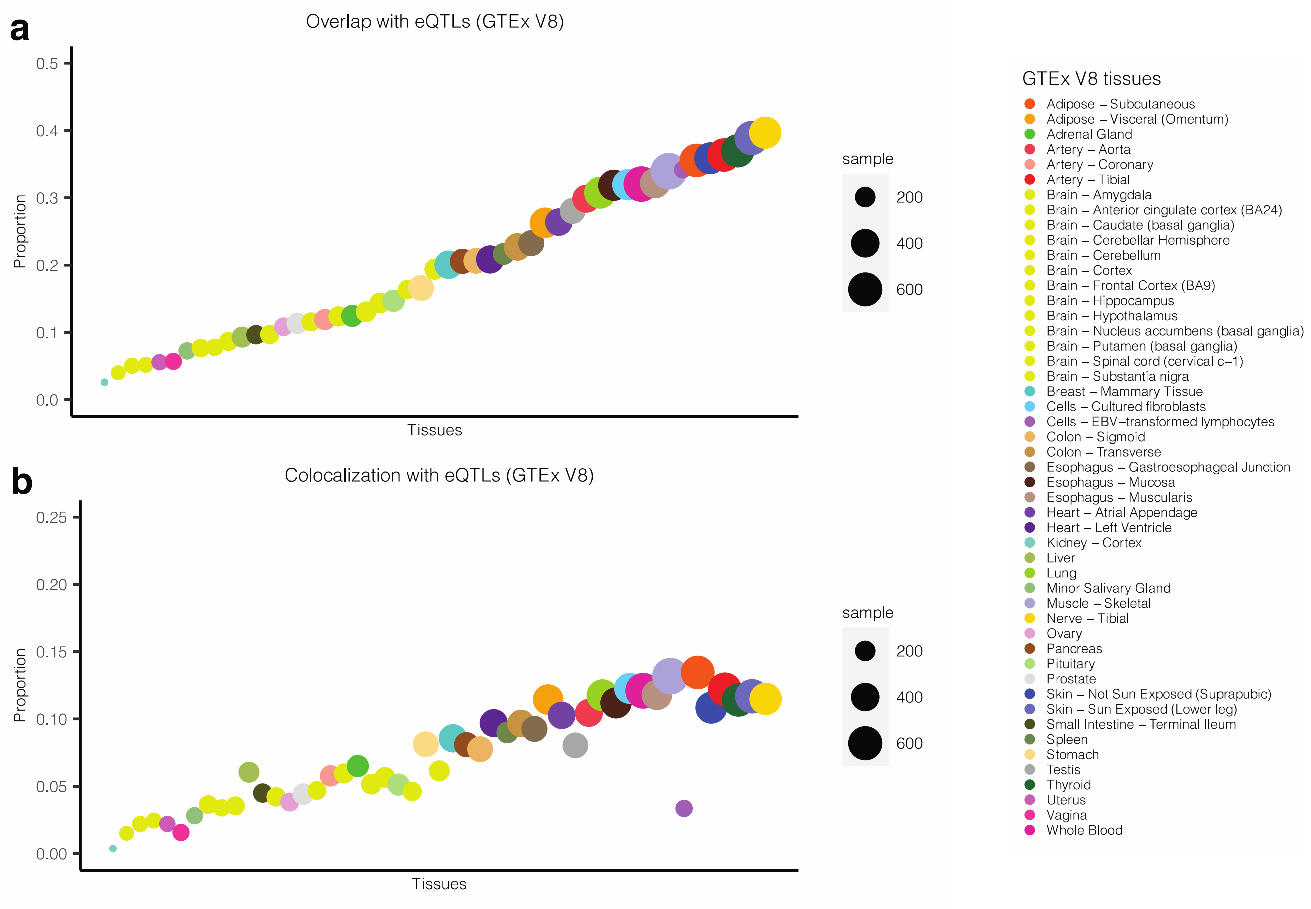


Extended Data Fig. 2

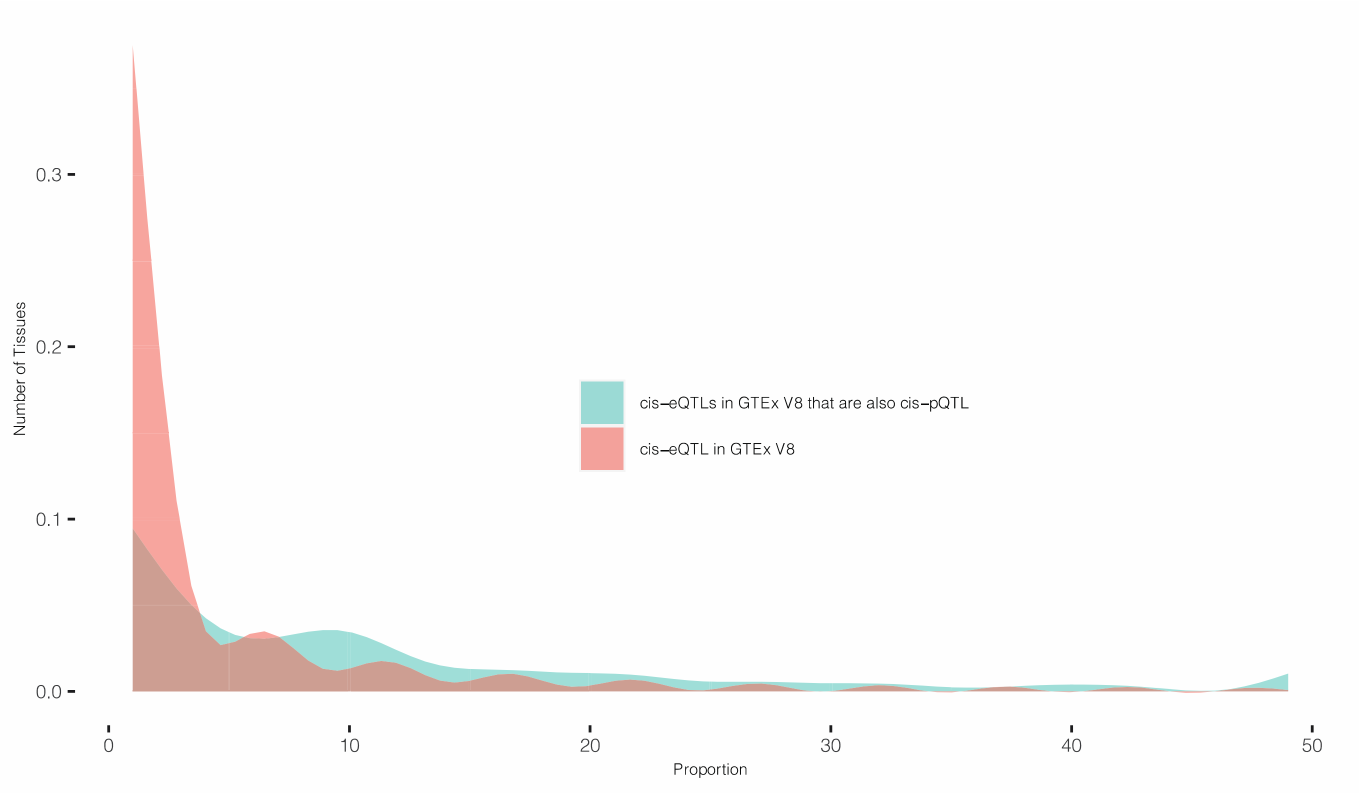


Extended Data Fig. 3

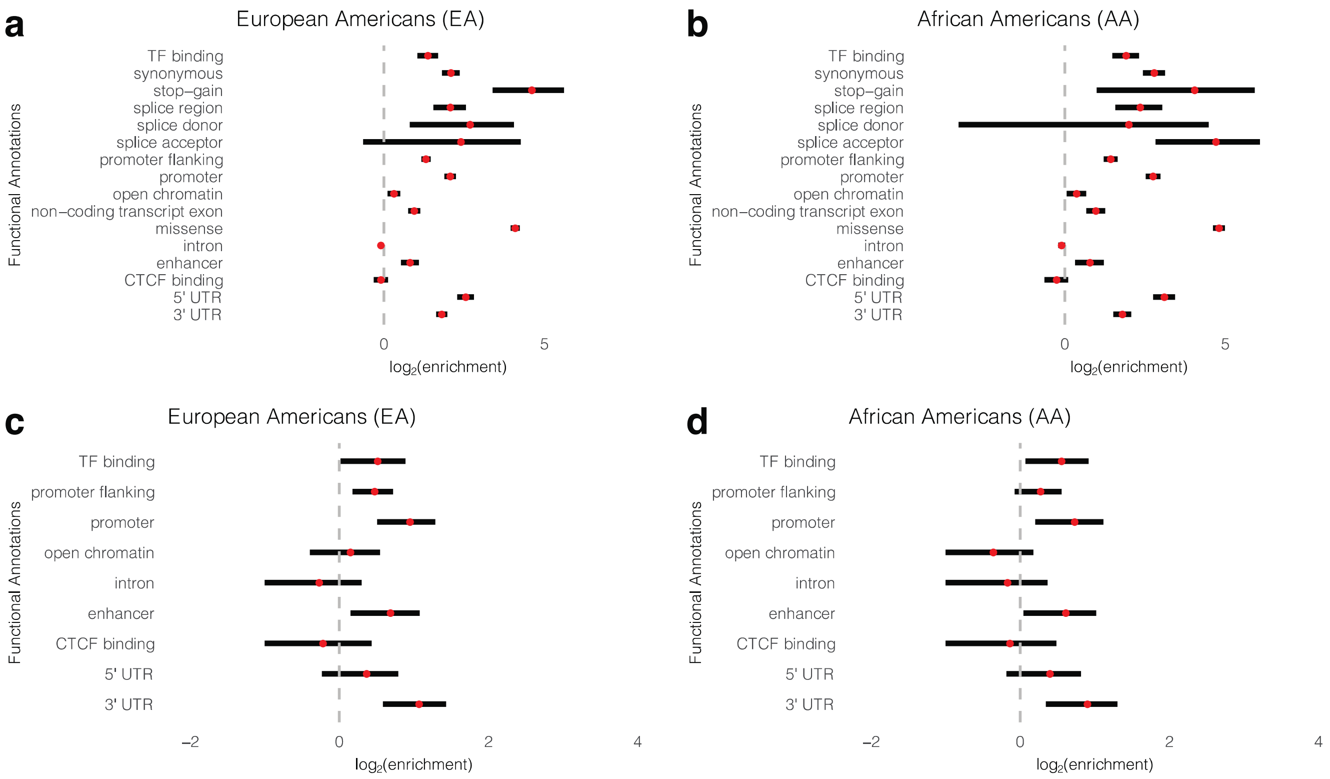


Extended Data Fig. 4

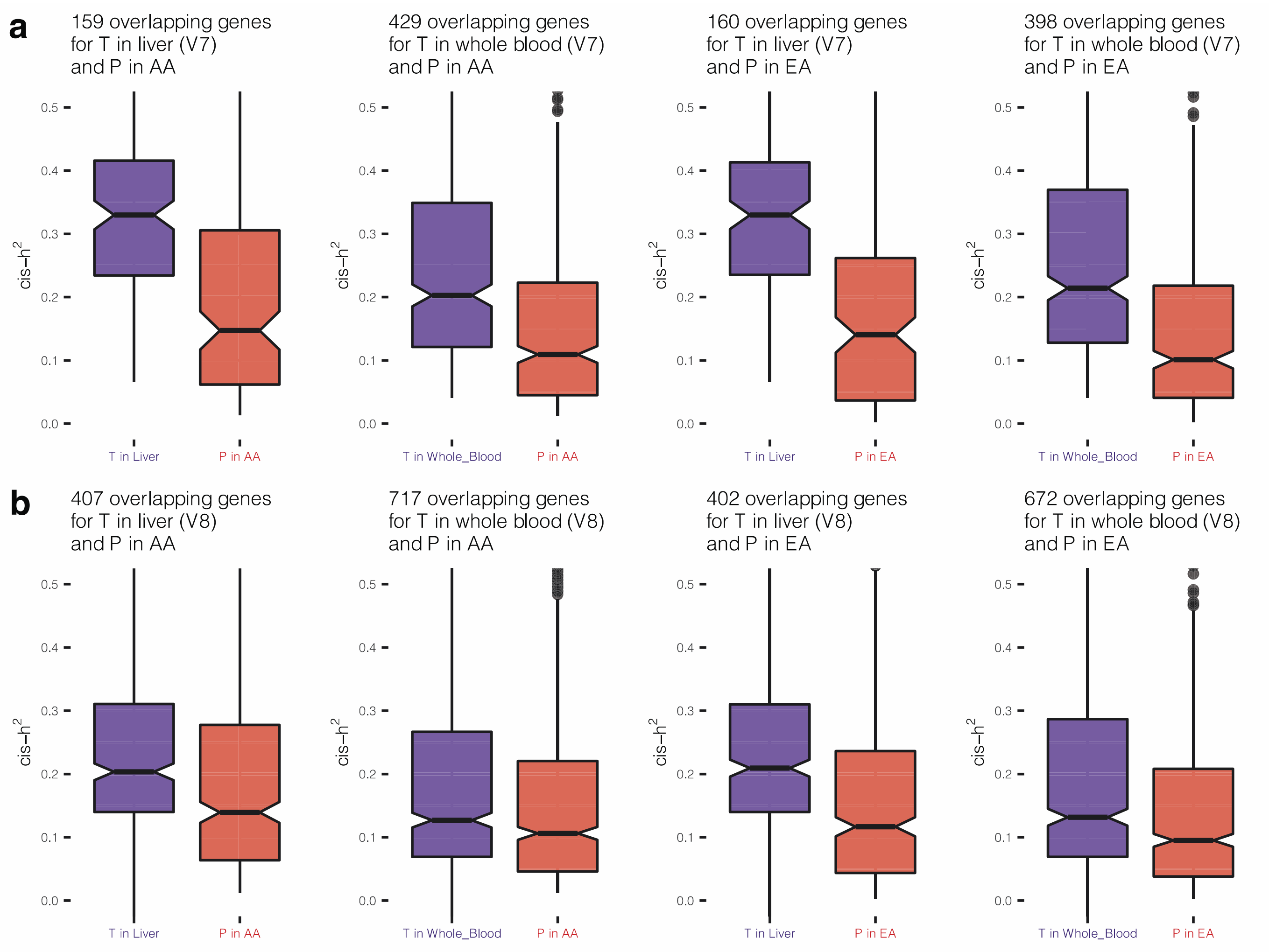


Extended Data Fig. 5

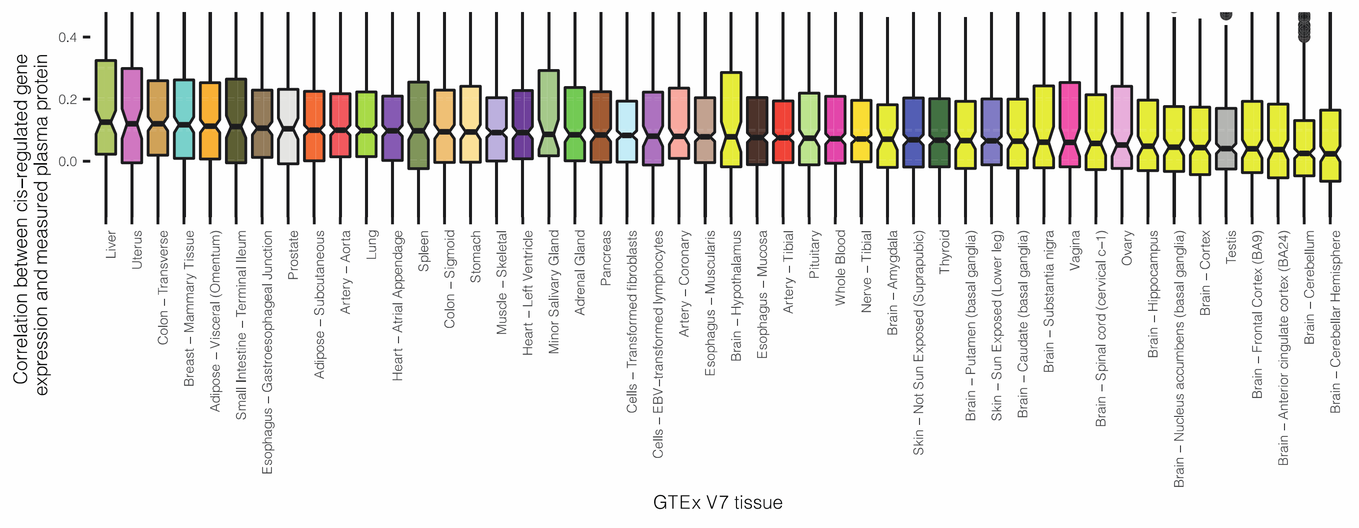


Extended Data Fig. 6

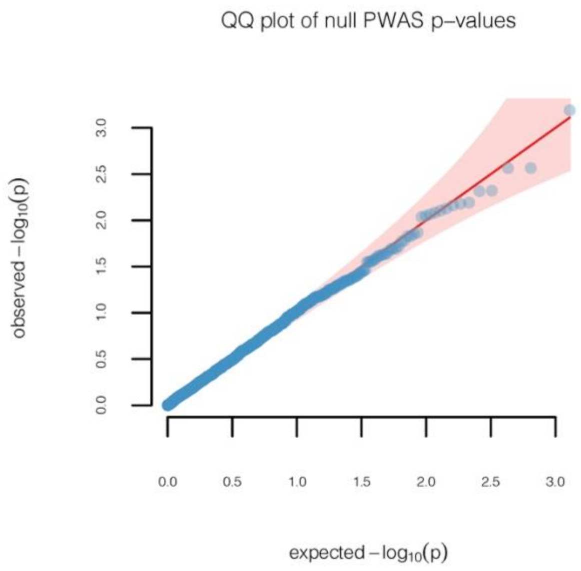


Extended Data Fig. 7

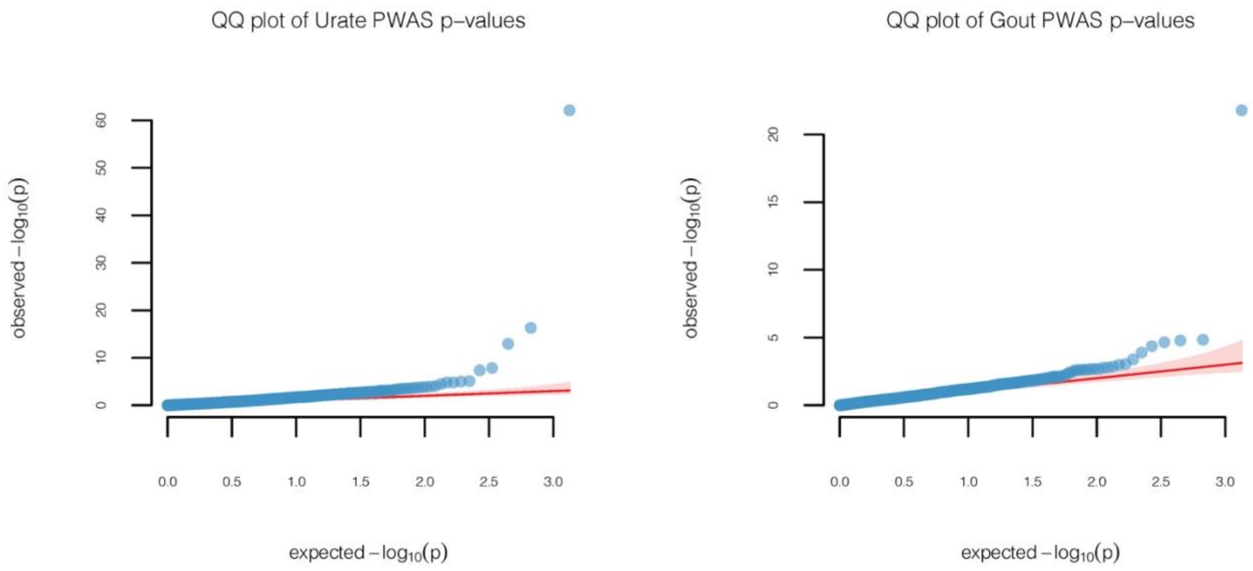


Extended Data Fig. 8

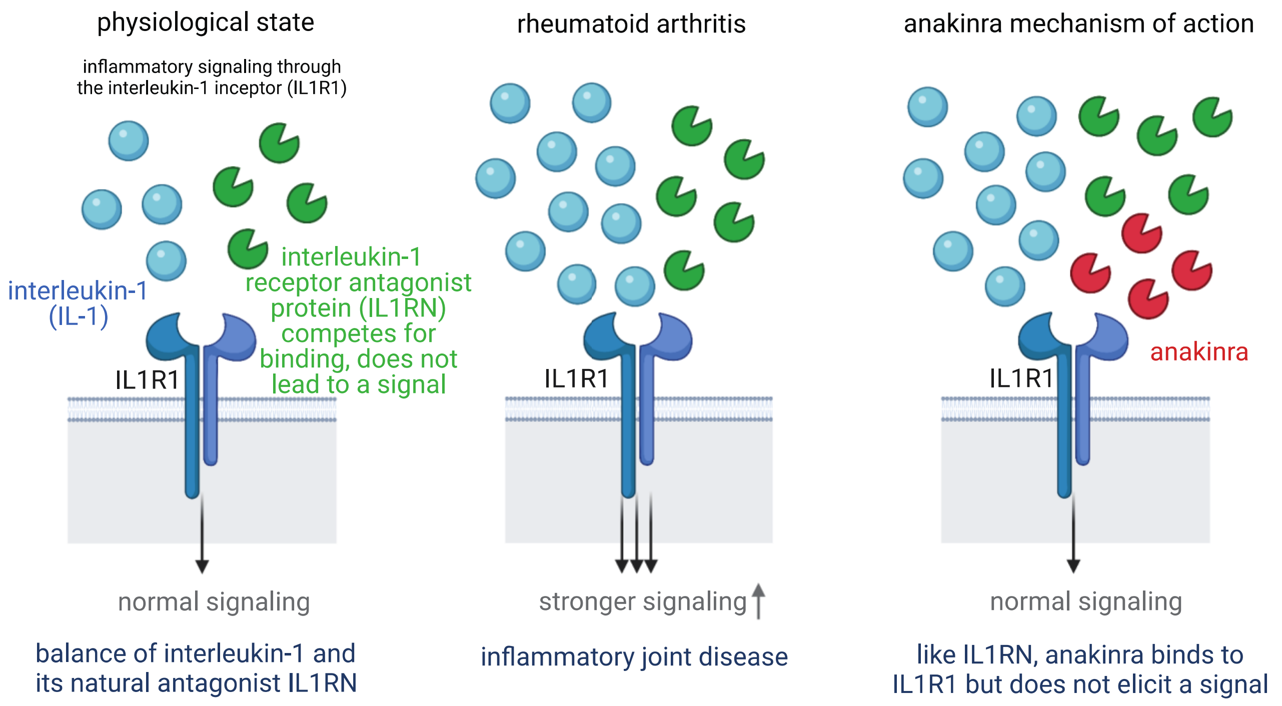


1. Anakinra is a synthetic drug that mimics the function of the natural protein IL1RN.
   It is approved for treating rheumatoid arthritis.
2. Our study shows that genetically higher IL1RN levels show protection from gout.
3. This suggests that anakinra may also be effective to treat gout (repurposing).

Extended Data Fig. 9

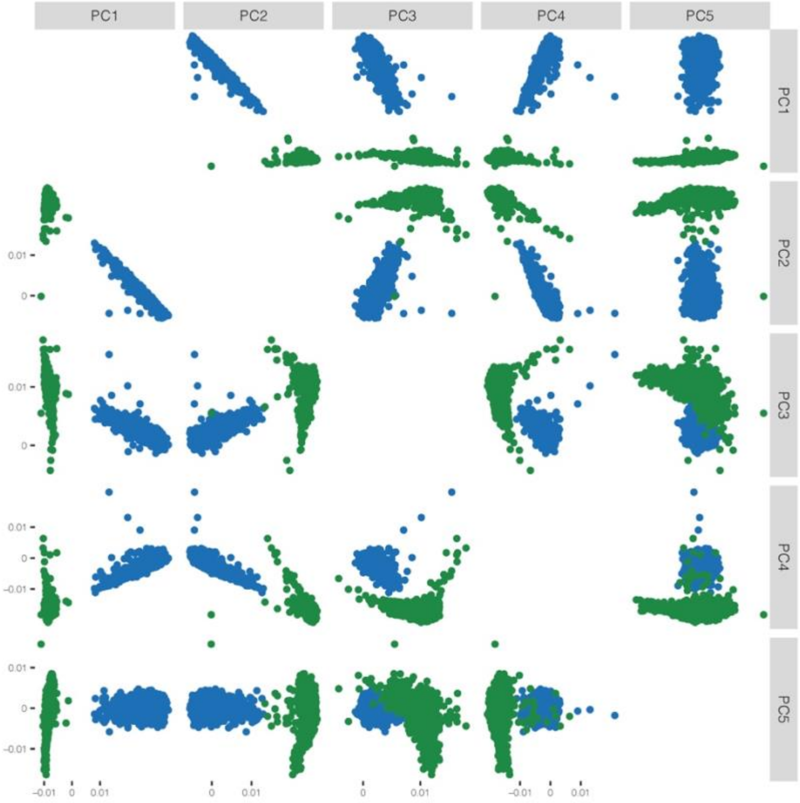


Extended Data Fig. 10
